# CD38 expression by neonatal human naïve CD4^+^ T cells shapes their distinct metabolic state and tolerogenic potential

**DOI:** 10.1101/2025.01.02.631038

**Authors:** Laura R. Dwyer, Andrea M. DeRogatis, Sean Clancy, Victoire Gouirand, Charles Chien, Elizabeth E. Rogers, Scott P. Oltman, Laura L. Jelliffe-Pawlowski, Susan V. Lynch, Rachel L. Rutishauser, Allon Wagner, Alexis J. Combes, Tiffany C. Scharschmidt

**Author notes:** Lead contact: Tiffany C. Scharschmidt. These authors contributed equally to this work (Andrea M. DeRogatis, Sean Clancy).

## Abstract

Neonatal life is marked by rapid antigen exposure, necessitating establishment of peripheral immune tolerance via conversion of naïve CD4^+^ T cells into regulatory T cells (Tregs). Here, we demonstrate heighted capacity for FOXP3 expression and tolerogenic function among cord blood versus adult blood naive CD4^+^ T cells and that this is linked to their unique metabolic profile and elevated expression of the NADase, CD38. Early life naïve CD4^+^ T cells demonstrate a metabolic preference for glycolysis, which directly facilitates their differentiation trajectory. We reveal an age-dependent gradient in CD38 levels on naïve CD4^+^ T cells and show that high CD38 expression contributes to both the glycolytic state and tolerogenic potential of neonatal CD4^+^ T cells, effects that are mediated at least in part via the NAD-dependent deacetylase SIRT1. Thus, the early life window for peripheral tolerance in humans is critically enabled by the immunometabolic state of the naïve CD4^+^ compartment.

## Introduction

Neonatal life is marked by rapid exposure of naïve immune cells to an array of new antigens, both self-antigens in peripheral tissues as well as exogenous microbial and environmental antigens. Developing adaptive immune tolerance to these antigens is critical to avoid excessive inflammatory or allergic responses that would be detrimental to homeostasis in the developing tissues and overall growth^1, 2^. The main cells responsible for establishment and maintenance of tolerance are regulatory T (Treg) cells, which express the transcription factor forkhead box protein 3 (FOXP3)^3, 4^. Tregs can use an array of effector molecules to suppress inflammation and maintain tolerance, including but not limited to surface molecules, such as CTLA4, CD25, and CD73, and immunosuppressives cytokine, such as IL-10 and TGF-β^5^. Notably, human fetal naïve CD4^+^ T cells have been shown to have a heightened capacity to differentiate into Tregs compared to their adult counterparts^5, 6^. Prior work has revealed higher expression of certain Treg-associated genes in human naïve CD4^+^ T cells during fetal development^7^, with a progressive decline of this signature in cord blood (CB) and adult blood (AB) naïve CD4^+^ T cells^8^, which may contribute to age-dependent differences in their function. Extension of heightened Treg generation into the post-natal period have been demonstrated in mice^9, 10^. In humans, the heightened prevalence of Tregs in pediatric tissues^11^ and the capacity to prevent certain food allergies via dietary exposure in infancy^12^ also hint at an early life window of tolerance. Indeed, prior work has indicated that CB CD4^+^ T cells may have a higher propensity to upregulate Treg markers and manifest suppressive capacity^13^. However, much remains undefined regarding the potential increased capacity for Treg differentiation among neonatal human T cells and what mechanisms might support this.

Metabolic pathways influence immune cell fate and function by directly modulating substrates needed for energy, protein and nucleotide production. They also impart key effects on cellular responses, for example through the role of nicotinamide adenine dinucleotide (NAD+) as a co-factor for the sirtuin family of histone deacetylases^14^ or direct modifications of histones by lactate^15^. Thus far, most work has focused on examining the metabolic state of adult immune cells, with comparatively little attention paid to how this might differ in early life. Even so, some key overarching principles have emerged that can help inform hypotheses about the metabolism of early life naïve CD4^+^ T cells^16^. Naïve cells are relatively quiescent and their low ATP needs are fulfilled primarily by mitochondrial oxidative phosphorylation (OXPHOS). With activation, T cells meet their heightened energy needs by rapidly increasing both OXPHOS and glycolysis, but their ATP dependency shifts towards glycolysis as a more immediate energy source. Finally, the transition into a memory state is paired with reduced glycolysis and a shift back to OXPHOS for energy generation^17, 18^. One of the few studies to explore early life metabolism focused on neonatal CD8^+^ T cells, identifying that an early life preference for glycolysis likely contributes to their preferential effector differentiation while limiting their ability to become memory cells^19^. Analogous studies to examine the metabolic state of neonatal naïve CD4^+^ T cells have yet to be performed, but are particularly important given the multiple differentiation paths open to naïve CD4^+^ T cells, i.e. differentiation into Tregs or various helper T cell types such as a Th1, Th2, or Th17, and the long-term impact these early cell fate decisions have on overall adaptive immune function and tolerance.

CD38, also known as cyclic ADP ribose hydrolase, is a glycoprotein expressed both intra-and extracellularly on many cell types, including immune cells^20^. CD38 has been used as a marker of early T cell activation^21^ as well as a marker for CD8^+^ T cell exhaustion or metabolic dysfunction in the setting of chronic viral infection^22^ or cancer^23, 24^. It was also recently reported as a potential marker of recent thymic emigrants (RTEs) in humans^25^. CD38 has various cellular functions, among the most prominent of which is regulating NAD+ levels by catalyzing the synthesis of ADP ribose and cyclic ADP ribose (cADPR) from NAD+, depleting NAD+ levels^20, 26^. CD38 also augments cellular calcium through cADPR’s role as an endogenous Ca2+ mobilizing nucleotide^27^ and facilitates tyrosine kinase activity downstream of TCR signaling via direct interaction of its cytoplasmic tail with the Lck SH2 domain^28^. Thus far, a few reports have connected CD38 levels to the metabolic state of adult T cells. In murine CD4^+^ T cells, deletion of CD38 increased OXPHOS capacity^23^. Similarly, in human adult naïve CD4^+^ T cells, pharmacological inhibition of CD38 via the compound 78c increased OXPHOS and mitochondrial mass^29^. Separately, one study computationally predicted that CD38 is positively associated with a pro-tolerance phenotype in murine Th17 cells^30^. Thus, existing evidence links CD38 expression to reduced OXPHOS dependence with potential implications for CD4^+^ T cell function. However, its contribution to the distinct metabolic state and differentiation capacity of neonatal T cells has not been examined. CD38’s enzymatic reduction of NAD+ levels, i.e. its NADase activity, has the potential to impact the metabolic and functional states of immune cells through altered activity of NAD-dependent acetylases, which can modulate chromatin structure or protein degradation^31^. Indeed, sirtuin 1 (SIRT1) is a prominent member of this NAD-dependent deacetylase family, which has been shown to shape mitochondrial biogenesis, influence autoimmunity, and even facilitate FOXP3 protein degradation^14, 32–35^. This link offers a potential mechanism of how CD38 could impact the functional and metabolic state of early-life immune cells that merits further study.

Here, we aimed to build on and interconnect knowledge in these areas to define how the metabolic state of early life human naïve CD4^+^ T cells shapes their capacity to support age-specific immune functions. We find that the unique metabolic state of neonatal naïve CD4^+^ T cells, i.e. their glycolytic preference, is intricately tied to high CD38 expression, and that both these age-specific features support their enhanced capacity for FOXP3 expression and tolerogenic capacity. We further identify CD38’s NADase function and downstream effects on SIRT1 as a central pathway mediating CD38’s impact on neonatal CD4^+^ T cell biology. Collectively, this data offers novel insight into a key time for immune education and function, and a new mechanism by which the early life window of tolerance may be reinforced in humans.

## Results

### CD38 is highly expressed on naïve CD4^+^ T cells in early life

To investigate key features distinguishing the neonatal immune system, we first performed flow cytometry on cord blood (CB) and adult blood (AB) peripheral blood mononuclear cells (PBMCs) using a panel incorporating key markers of cell identity and state. Immediately striking from these data was the observation that CD3^+^ CB T cells expressed high levels of extracellular CD38 compared to their adult counterparts (**Figure 1A, S1A**). Further sub-gating revealed the highest CD38 expression was on CB CD4^+^ T cells, with slightly less on CB CD8^+^ T cells and even lower CD38 levels on B cells or monocytes (**Figure 1B**). CD38 expression was notably highest on CB naïve CD4^+^ T cells followed by Tregs, with CB naïve CD4^+^ T cells exhibiting on average 4 to 5 fold more CD38 surface protein versus those in AB (**Figure 1C**). This result was intriguing given the established role of CD38 in cellular metabolism^20, 21^ and prior work suggesting a strong impact of metabolic state on the distinct differentiation potential of early life lymphocytes^36, 37^.

**Figure 1:**
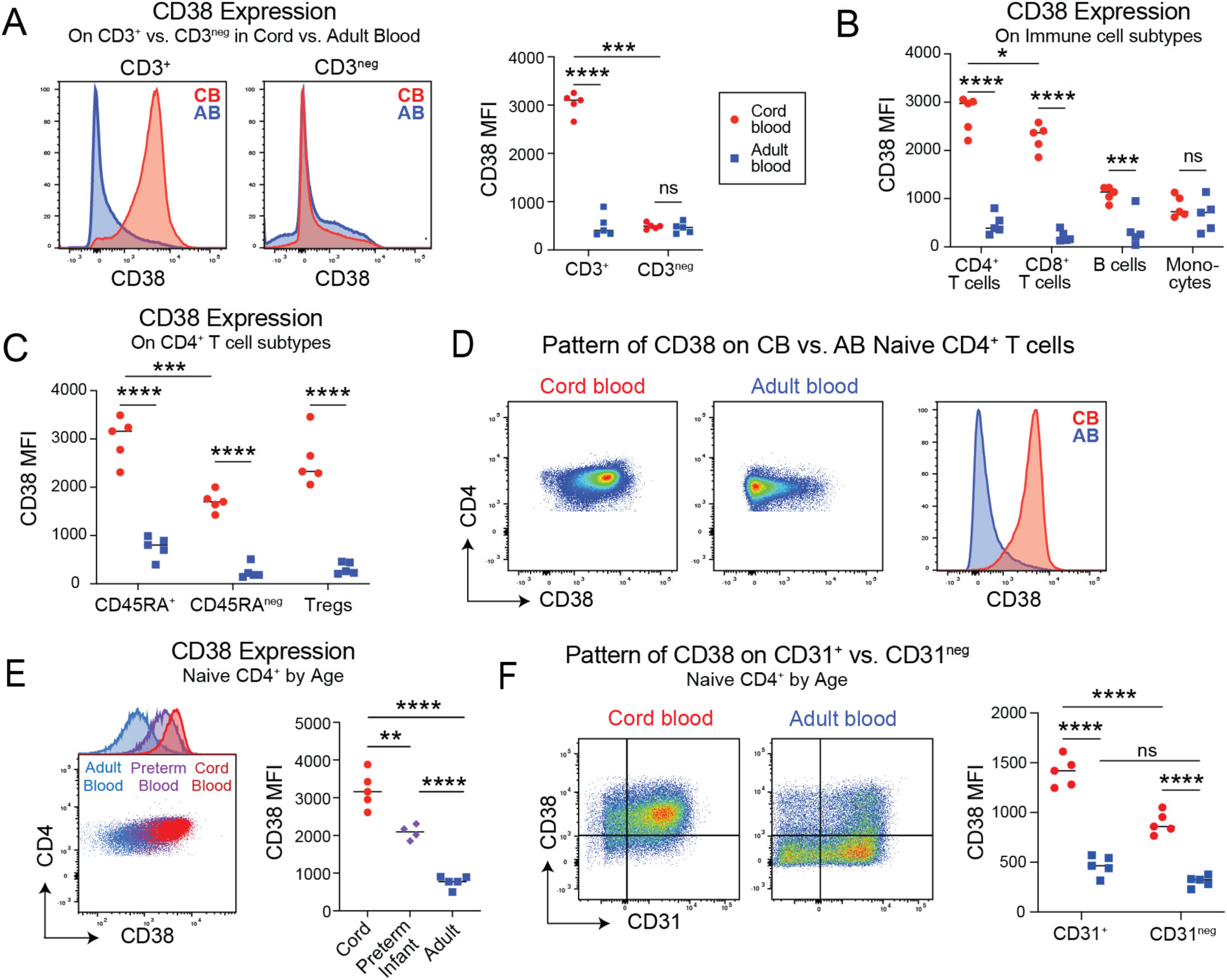
CD38 expression is highly enriched on naïve CD4^+^ T cells in early life. (A) Mean fluorescent intensity (MFI) of surface CD38 measured by flow cytometry on CD3 positive and CD3 negative cells from non-activated peripheral cord blood (CB) and adult blood (AB). (B) MFI of surface CD38 measured by flow cytometry on various immune cell populations (CD4^+^ T cells, CD8^+^ T cells, B cells, and monocytes) in non-activated peripheral CB and AB. (C) CD38 MFI on CD4^+^ T cell subsets (CD45RA^+^, CD45RA^neg^ and FOXP3^+^CD25^+^ Tregs) in non-activated peripheral CB and AB. (D) CD38 MFI on naïve CD4^+^ T cells from CB and AB. (E) CD38 MFI on naïve CD4^+^ T cells from CB, peripheral blood from 6 week-old preterm infants, or AB. (F) CD38 and CD31 expression on non-activated naïve CD4^+^ T cells from CB and AB and the CD38 expression pattern on CD31^+^ vs CD31^-^ cells by age. B, C, D, and F show one representative experiment from two experimental replicates. E shows one experiment.

Examination of CD38 staining on CB versus AB naïve CD4^+^ T cells revealed a largely unimodal pattern for each population with minimal overlap in expression level between ages (**Figure 1D**). Combined intracellular with extracellular staining for CD38 preserved the large age-dependent differences, confirming that the increase in CB expression was not due to internalization of CD38 in AB cells but instead overall expression (**Figure S1B**). As CD38 is often used as a marker of T cell activation^38^, we performed longitudinal measurements of its expression on CB or AB naïve CD4^+^ T cells that were isolated via negative selection (EasySep^TM^) then cultured in vitro in the presence of human anti-CD3 and anti-CD28 soluble antibody complexes (ImmunoCult^TM^). This confirmed an initial increase in CD38 expression in CB naïve CD4^+^ T cells after 3 hours of stimulation, with a similar but smaller, non-significant increase in AB at this same timepoint (**Figure S1C**). Of note, despite equivalent TCRβ expression on CB and AB naïve CD4^+^ T cells (**Figure S1D**), we did observe higher Nur77 staining at 3 hours post-stimulation (**Figure S1E**), consistent with prior literature indicating higher TCR reactivity of early life T cells^39^. Preservation of age-dependent differences in CD38 expression were maintained through 48 hours of stimulation, after which CD38 expression in CB declined to levels seen in AB naïve CD4^+^ T cells (**Figure S1C**).

To understand whether elevated CD38 expression reflects a unique CB state or a broader feature of the early life window, we performed CD38 staining on additional prenatal and postnatal samples. Staining of second trimester fetal spleen in parallel with CB and AB revealed even higher median CD38 expression on fetal naïve CD4^+^ T cells (**Figure S1F**). We also obtained peripheral blood from prematurely born infants aged 6 weeks (Adjusted Gestational Age: 0-1 weeks). CD38 staining showed intermediate levels on neonatal naïve CD4^+^ T cells as compared to CB and AB cells (**Figure 1E**). Taken together with the fetal data, this suggests an age-dependent gradient of CD38 expression. Based on the high rate of thymic output in early life and recent work showing that CD38 is elevated among recent thymic emigrants (RTEs)^40^, we performed CD38 staining on CB and AB naïve CD4^+^ T cells in parallel with the RTE marker CD31^41^. This confirmed higher CD38 expression on CD31^+^ versus CD31^neg^ cells in CB, with a non-significant trend towards higher CD38 in CD31^+^ versus CD31^neg^ in AB. However, large age dependent differences persisted both among CD31^+^ and CD31^neg^ subsets (**Figure 1F**). Thus, while CD38 levels do positively correlate with CD31 as a marker of RTE status, elevated CD38 expression on early life naïve CD4^+^ T cells is not merely a reflection of RTE frequency^18^.

### Neonatal versus adult naïve CD4^+^ T cells demonstrate a distinct transcriptional trajectory following stimulation

To further elucidate key features distinguishing CB versus AB naïve CD4^+^ T cells, the distinct functional potential of these cells, and the relationship to CD38 expression, we designed a longitudinal experiment to transcriptionally profile by RNA sequencing (RNA-seq) CB versus AB naïve CD4^+^ T cells at baseline and three time points following activation (**Figure S2A**). Naïve CD4^+^ T cells (CD4^+^ CD8^neg^ CD45RA^+^ CD27^+^ CCR7^+^ CD95^neg^) were isolated by FACS from CB and AB samples (**Figure S2B**). Cells were then either immediately harvested (0 hour) or first stimulated in culture with anti-CD3 and anti-CD28 soluble antibody complexes for 3, 24, or 48 hours. RNA-seq was then performed on these bulk cell populations, generating a dataset that could be mined for both baseline transcriptional differences and distinct age-dependent trajectories post-activation.

Principal component analysis (PCA) was performed to visualize transcriptional differences between samples by time point and age (**Figure 2A**). PC1 encompassed 47.9% of variance and stratified samples longitudinally by time point, based on genes involved in cell proliferation and activation, e.g. *CDC25A*, *MCM10*, *MKI67*, *IL2* (**Figure S2C**). PC2 13.2% variance separated CB from AB naïve CD4^+^ T cells, with several of the axis-driving gene expression features having been previously reported to be differentially expressed by age or associated with development, e.g. *SOX4*^42^, *BCL11A*^43^, *TSHZ2*^7^ (**Figure S2D, S2E**). A heatmap generated on the most variable features revealed gene clusters with distinct patterns of expression based on CB versus AB status or time point (**Figure 2B**). Some clusters encompassed genes that varied between CB and AB, largely independent of activation status (2, 6, 8) whereas others showed dynamic increases (4, 5) or decreases (3, 7) in expression levels post-stimulation (**Figure 2C**). For example, Cluster 4 contained several cytokine genes that were lowly expressed at baseline or 3 hours but had increased in either AB or CB by 24 to 48 hours. Specifically, whereas AB naïve CD4^+^ T cells upregulated IFNG, those from CB showed increased expression of *IL4*, *IL10*, and *IL13*. Notably, differential expression analysis at each time point revealed enrichment of key tolerance-related genes in CB versus AB. At previously reported^8^, the gene encoding Helios, *IKZF2*, was already elevated in resting CB naïve CD4^+^ T cells. Other tolerance-associated genes increased in CB only after stimulation, e.g. *IL2RA*, *IL10*, *GATA3*, *PPARG*, and *PDCD1*. Conversely, certain CD4^+^ T effector genes were preferentially increased in AB with stimulation, i.e. *IL2* and *IFNG* (**Figure 2D**). Gene set enrichment analysis (GSEA)^44, 45^ of ranked genes in resting naïve CD4^+^ T cells showed AB enrichment in host defense and cytokine responses, with more subtle enrichment of development-related pathways in resting CB (**Figure S2F**). After 48 hours of stimulation, this developmental signature in CB was accompanied by enrichment for processes such as regulation of cell activation, adhesion and proliferation. Stimulated AB was instead enriched at 48 hours for terms related to defense against symbionts or virus, positive regulation of pattern recognition receptor signaling, and IL-17 production (**Figure S2G**). Taken together, this analysis revealed a distinct transcriptional signature for CB cells both at baseline and up to 48 hours following activation, with features suggestive of a pro-tolerogenic signature in CB.

**Figure 2:**
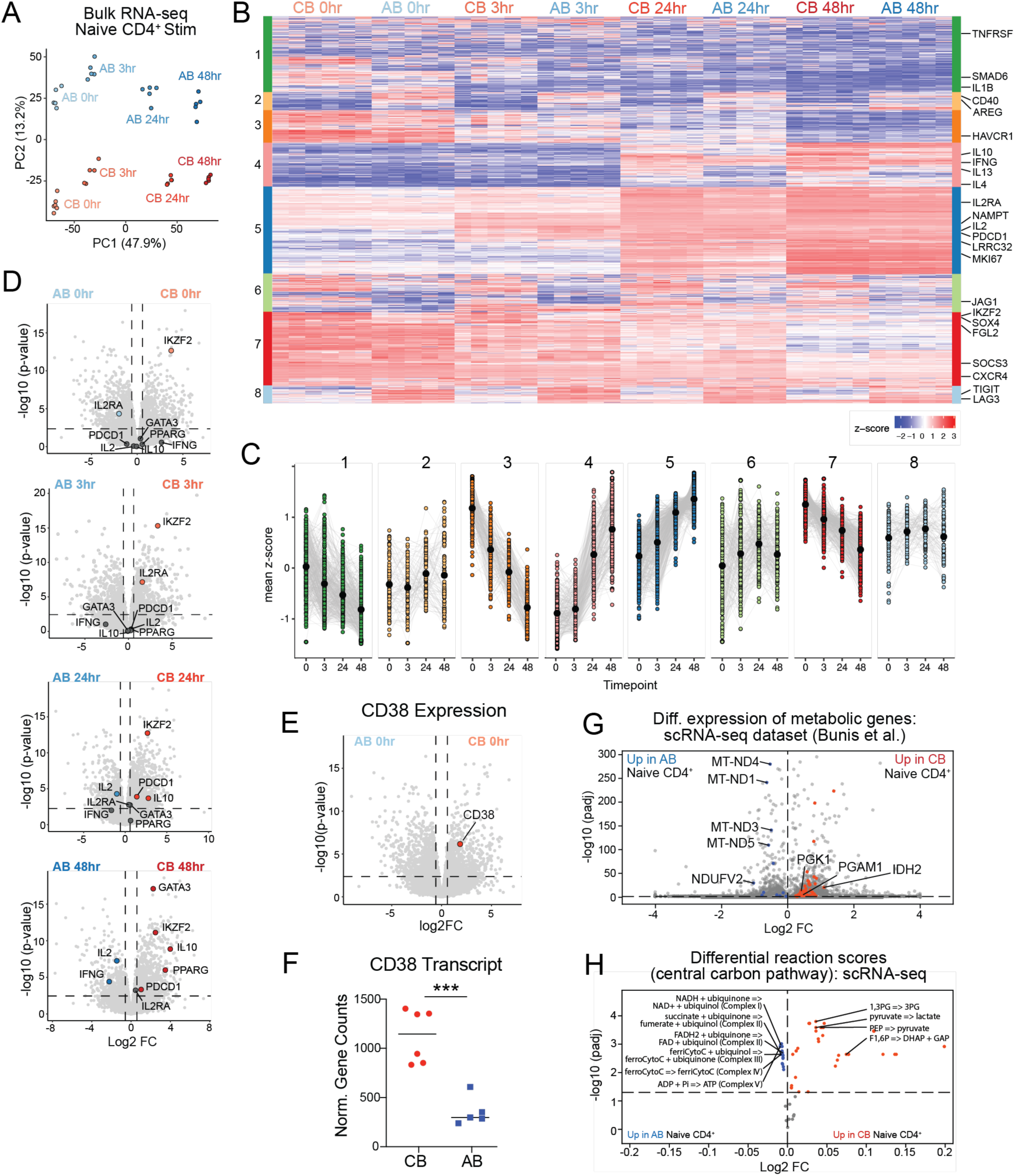
Transcriptomic data analysis and in-silico modeling of CB versus AB naïve CD4^+^ T cells reveals unique functional and metabolic programs at baseline and after activation. (A) PCA plot of bulk RNA-seq data from naïve CD4^+^ T cells (CD4^+^CD8^neg^CD45RA^+^CD27^+^CCR7^+^CD95^neg^) sorted from CB and AB, with samples taken at baseline (0 hr), 3, 24 and 48 hr after stimulation. (B) Heatmap of top features (i.e. clusters of genes) with variable expression by age (CB versus AB), each cluster is denoted by a different color on x axis. (C) Mapping of cluster expression (mean z-score of each gene) over time. Black dots represent the mean z-score per time point. Colors correspond with labelled clusters in B. (D) Volcano plots of differential gene expression for CB versus AB naïve CD4^+^ T cells at each time point, with specific genes related to tolerance or activated effector cells highlighted. Horizontal dotted line represents Benjamini-Hochberg adjusted p-value of 0.05. (E, F) Volcano plot highlighting CD38 from RNA-seq of CB versus AB naive CD4^+^ T cells (non-activated = 0hr) and normalized gene counts of CD38. (G) Differential expression of key metabolic genes from re-analysis of published scRNA-seq dataset of CB versus AB FASC-sorted naïve CD4^+^ T cells (Bunis et al.). (H) Compass comparison of CB versus AB naïve CD4+ T cell metabolic states in central carbon metabolism reveals upregulation of core glycolysis reactions and an overall downregulation of the OXPHOS pathway in CB. Dots represent metabolic reactions, as annotated in the Human1 GEM metabolic model.

### The transcriptome of neonatal naïve CD4^+^ T cells suggests a distinct metabolic profile

Based on existing literature linking the metabolic state of neonatal naïve CD8^+^ T cells to their differentiation trajectory and functional capacity^19, 46^, we next wanted to investigate if there were baseline differences in the metabolic state of CB and AB naïve CD4^+^ T cells. Focused examination of our bulk RNA-seq dataset for differentially expressed metabolic genes, revealed roughly 4-fold higher levels of *CD38* transcript in CB versus AB naïve CD4^+^ T cells, corroborating our flow cytometry data (**Figure 2E-F**). Certain glycolysis-related genes were also elevated in CB at baseline (**Figure S2H**), hinting that neonatal naïve CD4^+^ T cells display a distinct metabolic profile. To corroborate this finding, we turned to an independent dataset previously published by Bunis et al., in which sorted CB and AB naïve CD4^+^ T cells were subjected to single-cell RNA sequencing (scRNA-seq)^8^. Differential gene expression analysis validated our observation of upregulation of glycolysis-related genes in CB and revealed a parallel downregulation of mitochondrial genes critical to oxidative phosphorylation (**Figure 2G**).

To further characterize the metabolic profile of CB and AB blood naïve CD4^+^ T cells, we turned to Compass, a flux-balance analysis algorithm that explicitly models the flow of metabolites through the metabolic network to characterize the metabolic state of cells^30^. Compass analysis of our own baseline bulk RNA-seq dataset did not reveal statistically significant pathways in CB versus AB. Thus, we leveraged the Bunis et al. scRNA-seq dataset, performing a Compass run on the central carbon pathways (glycolysis, TCA, and OXPHOS). To robustly capture the transcriptomic state of cells in face of inter-donor variation, we pseudobulked (averaged), the gene expression profiles of cells belonging to the same donor. Using these pseudobulk profiles, we discovered striking metabolic differences where glycolysis reactions were upregulated, while OXPHOS reactions across all complexes were downregulated in the CB group (**Figure 2H**). Thus, our metabolic-focused analyses of the transcriptional activity of naïve CD4^+^ T cells suggest that CB cells have a distinct baseline metabolic state, with a possible preference for glycolysis.

### Neonatal naïve CD4^+^ T cells demonstrate more tolerogenic potential upon stimulation than their adult counterparts

We next sought to interrogate the functional relevance of the increased tolerance-related gene expression among stimulated CB naïve CD4^+^ T cells seen in our RNA-seq dataset. To replicate our RNA-seq conditions, we provided T cell receptor (TCR) stimulation without added cytokines. Specifically, thawed aliquots of CB and AB mononuclear cells were subject to a naïve CD4^+^ T cell negative selection kit (EasySep^TM^), which effectively isolated CD4^+^CD45RA^+^ T cells that were CD25^neg^ and FOXP3^neg^ (**Figure S3A-B**). These cells were then cultured in the presence of human anti-CD3 and anti-CD28 soluble antibodies (ImmunoCult^TM^) with no added cytokines, referred to as our “No Cytokine” condition. Separately, we performed analogous stimulation in the presence of IL-2 and TGFβ, replicating published conditions for “induced Treg” generation^7^, referred to as our “iTreg” or “IL-2+TGFβ” condition. Cells were harvested after 96 hours and analyzed by flow cytometry. Despite equivalent expression of CD69, a marker of activation, and Ki67, a marker of recent proliferation (**Figure S3C-D**), stimulated CB and AB CD4^+^ T cells demonstrated distinct patterns of FOXP3 and CD25 expression (**Figure 3A**). Under “No Cytokine” conditions, a higher percentage of CB versus AB cells became CD25 positive (**Figure 3B**). Irrespective of age, the majority of these CD25^+^ cells expressed FOXP3 at levels approaching that of endogenous blood Tregs (**Figure 3C, S3E**), thus we will refer to them hereafter as FOXP3^+^CD25^+^ cells. This heightened frequency of FOXP3^+^CD25^+^ cells from CB was replicated under “No Cytokine” conditions using various methods of CD3/CD28 stimulation (**Figure S3F**). In the presence of IL-2+TGFβ, the percentage of FOXP3^+^CD25^+^ cells again trended higher in CB versus AB but did not reach statistical significance (**Figure 3B**). As expected from iTreg culture conditions, these cells expressed much higher levels of FOXP3 than those generated without cytokines. Notably, however, iTregs generated from CB had higher FOXP3 levels than their AB-derived counterparts (**Figures 3A, 3C**).

**Figure 3:**
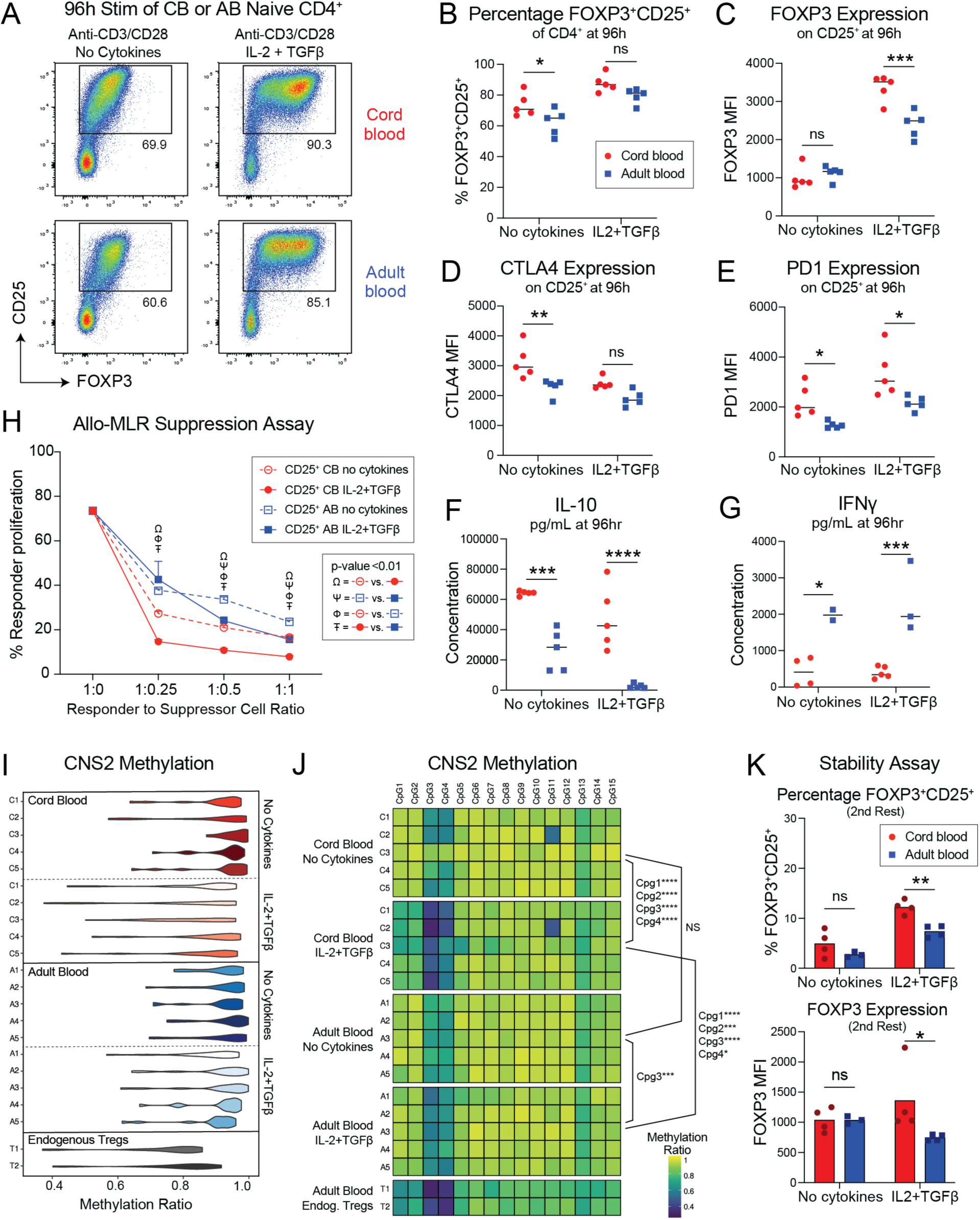
Neonatal naïve CD4^+^ T cells demonstrate more tolerogenic potential upon stimulation than their adult counterparts. (A,B) Representative flow plots and graph of the percentage of FOXP3^+^CD25^+^ cells from cord blood (CB) and adult blood (AB) naïve CD4^+^ T cells at 96hr following stimulation with Immunocult^TM^ anti-CD3/CD28 soluble antibodies with no added cytokines or with IL-2 (10 ng/ml; PeproTech) and TGF-β (50 ng/ml; PeproTech). (C-E) Expression of various markers on FOXP3^+^CD25^+^ at 96hr after CB or AB naïve CD4^+^ T cell stimulation, as measured by mean fluorescence intensity (MFI) by flow cytometry. (F,G) ELISA-based measurement of cytokines in cell culture supernatant of stimulated CB and AB naïve CD4^+^ T cells at 96hr. (H) Allogenic mixed lymphocyte reaction (Allo-MLR) of the suppressive capacity of CB and AB “No cytokine” and “iTreg” stimulated cells (see Figure S3N). (I, J) FOXP3 methylation was assessed in CB and AB where naïve CD4+ T cells were activated with Immunocult^TM^ anti-CD3/CD28 soluble antibodies and cultured for 96hr under no cytokine or IL-2+TGF-β conditions. Cells were then sorted to obtain CD4^+^CD25^+^CD127^LO^ cells. Additionally, CD4^+^CD25^+^CD127^LO^ cells were sorted from adult PBMCs to obtain endogenous Tregs. Sorted cells were then sent to Zymo Research to undergo targeted bisulfite sequencing of the CNS2 region of the FOXP3 locus to assess methylation patterns between ages and treatment conditions. Heatmap and violin plot of methylation ratios of CpG islands shown, statistically significant differences in methylation ratio for various CpG sites by age or culture condition via one-way Anova are shown in (J). (K) FOXP3 stability was assessed in CB and AB naïve CD4^+^ T cells through two rounds of stimulation and rest (see Figure S3P,Q). Data shown here is following the second rest. All data points represent individual samples from different biological donors. For A-E, and H, one representative experiment of two or three is shown. For F, G, I, J and K one experiment was done and is shown.

Deeper examination of FOXP3^+^CD25^+^ cells generated from CB versus AB revealed their higher relative expression of several Treg effector molecules. Specifically, they expressed more CTLA4, PD1, Helios or ICOS under either “No Cytokine” conditions, “iTreg” conditions, or both (**Figure 3D-E, S1G-H**). In contrast, similar levels of CD25 were seen on FOXP3^+^CD25^+^ cells irrespective of age or culture condition (**Figure S3I**). We additionally performed ELISA on culture supernatants at 96 hours post-stimulation to profile cell cytokine secretion. This revealed increased levels of IL-10 in CB supernatant under both “No cytokine” and “iTreg” conditions (**Figure 3E**). By comparison, AB culture supernatant contained more IFNγ under either condition (**Figure 3F**) and more IL-2 under “No Cytokine” conditions (**Figure S3J**), both of which are typically produced by CD4^+^ effector T cells. Aside from slightly more IL-4 in “No Cytokine” AB supernatant, no other significant age-dependent differences were seen for IL-4, IL-5 or IL-13 under either culture condition (**Figure S3K-M**). Thus, FOXP3^+^CD25^+^ cells generated from CB naïve CD4^+^ T cells exhibit elevated expression of various regulatory markers and a distinct, more tolerogenic, profile of cytokine secretion versus their AB-derived counterparts.

We next assessed the relative suppressive capacity of FOXP3^+^CD25^+^ cells derived from CB or AB under either “No Cytokine” or “iTreg conditions” in an allogeneic mixed lymphocyte reaction (MLR). Live CD4^+^CD25^+^CD127^Lo^ “Suppressor” cells were sorted from these cultures at 96 hours and co-cultured at varying ratios with CTV-labelled PBMC “Responder cells” from a single unrelated adult donor, and irradiated PBMC from a second, unrelated adult donor to serve the role of antigen presentation. The latter two populations were depleted of CD4^+^CD25^+^CD127^Lo^ via sorting prior to co-culture (**Figure S3N**). Flow-based analysis of “Responder cell” proliferation after 7 days revealed significant differences in the suppressive capacity of FOXP3^+^CD25^+^ cells, based on both age and culture conditions (**Figure S3O**). As anticipated, “iTreg” culture led to more potently suppressive cells versus “No Cytokine” stimulation for either age of naïve CD4^+^ T cell. Intriguingly though, CB-derived “Suppressor” cells were more potent than their AB-derived counterparts for both “No Cytokine” and “iTreg” conditions, demonstrating higher dose-dependent suppression of “Responder” cell proliferation at all 3 ratios tested (**Figure 3H**). Thus, FOXP3^+^CD25^+^ cells from CB express more suppressive capacity, especially those generated in the presence of IL-2 and TGFβ.

Treg stability and function has been linked epigenetically to demethylation of conserved non-coding sequence 2 (CNS2), a CpG-rich *FOXP3* intronic *cis-*regulatory element^47, 48^. We thus performed methylation analysis of this locus in sorted CD25^+^ cells generated from CB or AB under either “No Cytokine” or “iTreg” culture conditions. Inclusion of endogenous adult blood (CD4^+^CD25^+^CD127^Lo^) Tregs served as a positive control, revealing a methylation ratio ranging from 0.4 to 0.9 depending on the specific CpG site (**Figure 3I**). CD25^+^ cells derived from either CB or AB under “No Cytokine” conditions showed higher methylation ratios, ranging between 0.6 and 1, with no statistically significant differences seen by age. In contrast, culture with IL-2 and TGFβ led to significantly more demethylation among CB-derived CD25^+^ cells, specifically at CpG sites 1 through 4, when compared to either their CB “No Cytokine” or AB “iTreg” counterparts (**Figures 3I-J**).

Finally, we sought to assess if differential degrees of CNS2 demethylation among FOXP3^+^CD25^+^ cells generated from CB iTreg cultures corresponded to increased stability of FOXP3 expression. To do so, we adopted previously published methods for testing FOXP3 stability, in which cells are subjected to two sequential rounds of stimulation and rest, with flow-based assessment after each^49^. Specifically, we isolated naïve CD4^+^ T cells from CB or AB and stimulated them for 96 hours under either “No Cytokine” or “IL-2+ TGFβ” conditions. The cells were then washed and all groups rested with IL-2 prior to the “first rest” timepoint. Separate aliquots were then re-stimulated for 96 hours, this time will all groups also receiving IL-2, prior to another 96 hours of rest and assessment at the “second rest” timepoint (**Figure S3P**). At the “first rest” timepoint, no differences were seen between the ages or culture conditions with regard to either the percentage of FOXP3^+^CD25^+^ cells or their levels of FOXP3 expression (**Figure S3Q**). At the “second rest” timepoint, however, CB naïve CD4^+^ T cells initially subjected to iTreg conditions demonstrated significantly higher percentages of FOXP3^+^CD25^+^ cells than their AB counterparts, and these cells likewise expressed higher levels of FOXP3 protein (**Figure 3K**). Taken together, these experiments suggest that naïve CD4^+^ T cells from CB versus AB have a higher potential upon stimulation to become functionally suppressive cells, and that when cultured under “iTreg” conditions they demonstrate heightened Treg-associated properties, including suppression in allogeneic MLR, CNS2 demethylation and stability of FOXP3 expression following repeated rest and re-stimulation.

### Neonatal naïve CD4^+^ T cells express distinct metabolic molecules versus their adult counterparts

Based on prior work linking the metabolic state of neonatal murine naïve CD8^+^ T cells to their distinct differentiation trajectory^36, 50^, we wondered if metabolic features of CB naïve CD4^+^ T cells might promote their capacity to develop tolerogenic properties upon stimulation. The expression of metabolic proteins such as nutrient transporters, transcription factors, and metabolic enzymes measured via mass cytometry (CyTOF) strongly correlate with the metabolic state of T cells^51, 52^. Thus, we developed an immunometabolism-focused CyTOF panel (**Figure 4A**) and performed analysis of naïve CD4^+^ T cells (CD45RA^+^CCR7^+^) from five CB and five AB donor PBMCs to elucidate if distinct metabolic transcriptional signatures (**Figure 2G-H, S2H**) translated into age-related differences in metabolic function at the protein level. We first used multi-dimensional scaling (MDS) of median marker expression to examine the metabolic state of CB and AB naïve CD4^+^ T cells in an unsupervised manner. Distinct patterns of metabolic marker expression among CB and AB naïve CD4^+^ T was evident through their striking separation by age across MDS dimensions (**Figure 4B**). To examine the different metabolic states present in the naïve CD4^+^ T cell pool across age, we performed clustering with the FlowSOM^53^ algorithm using metabolic and reduced lineage markers which yielded 8 distinct clusters (**Figure 4C, S4A**). CB and AB naïve CD4^+^ T cells were largely distributed into distinct non-overlapping clusters, although each contained a small number of cells located in clusters predominantly associated with the other age group (**Figure 4D-E**).

**Figure 4:**
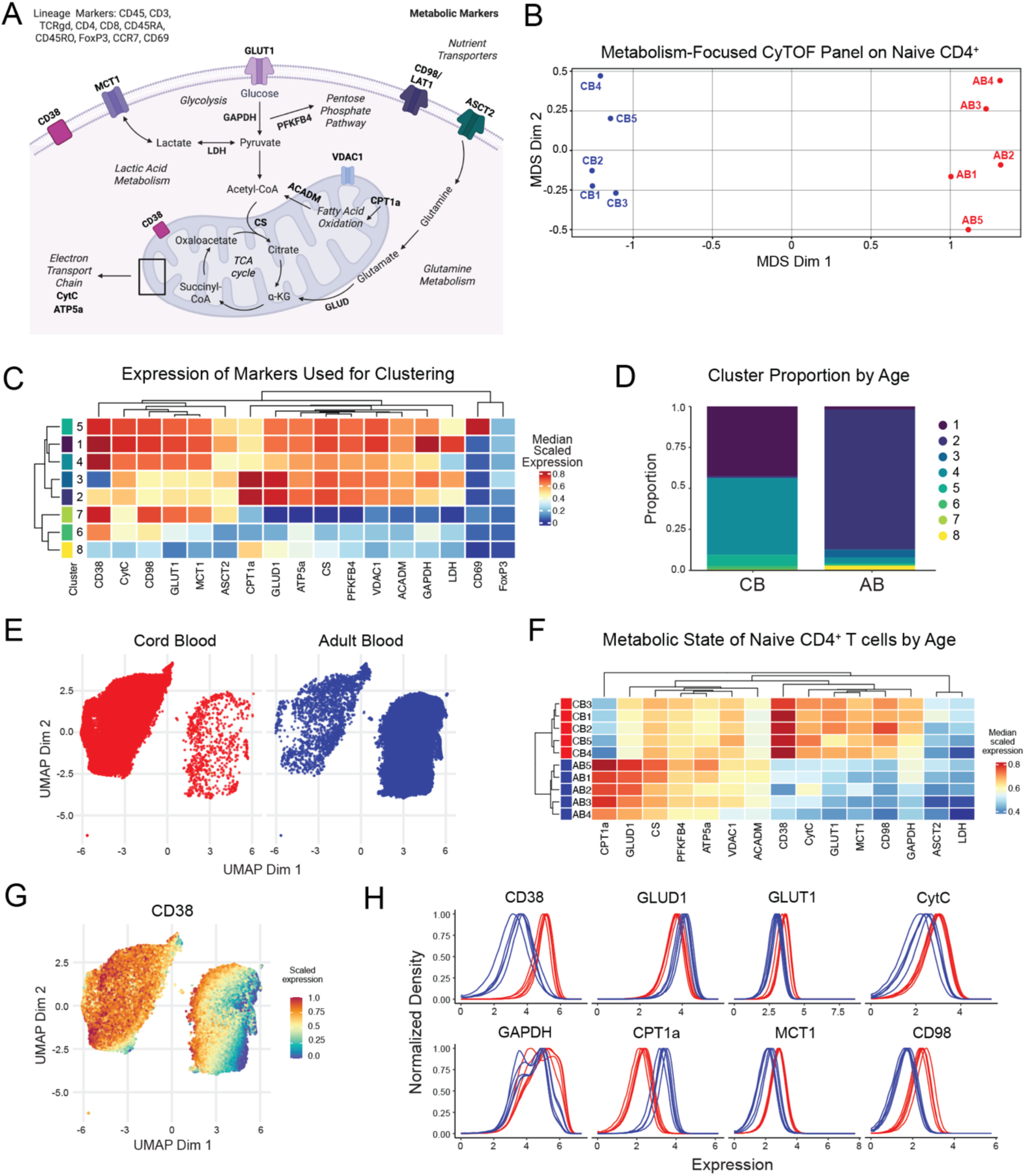
Neonatal naïve CD4^+^ T cells express distinct patterns of metabolic proteins versus their adult counterparts. (A) Schematic of markers used in metabolism-focused CyTOF panel. Metabolism markers are shown in bold in their corresponding pathways or functions. (B) Multi-dimensional scaling (MDS) of median marker expression was used to examine the similarity of CB and AB naïve CD4^+^ T cell metabolic state in an unsupervised manner. Markers used for MDS and clustering: CD98, CD69, LDH, PFKFB4, CS, ACADM, GLUD1, ATP5a, VDAC1, GLUT1, CytC, GAPDH, ASCT2, CPT1a, MCT1, CD38, FOXP3. (C, E) To examine the different metabolic states present in the naïve CD4^+^ T cell pool across ages, clustering was performed with the FlowSOM algorithm using metabolic and reduced lineage markers with CB and AB naïve CD4^+^ T cells separating into distinct clusters based on metabolic protein expression. (D) Differential abundance analysis showed CB naïve CD4^+^ T cells to be significantly enriched in clusters in clusters 1, 4 and 5. (F) Expression patterns of metabolic proteins across individual CB and AB naïve CD4^+^ T cells samples. (G) Intensity plot of CD38 expression in the UMAP space. (H) Expression patterns of metabolic markers for all ten CB and AB donors.

Consistent with our RNA-seq data, CB naïve CD4^+^ T cells showed increased expression of proteins central to glycolysis and decreased expression of molecules associated with fatty acid oxidation (FAO) (**Figure 4F**). Differential abundance analysis^54^ showed CB naïve CD4^+^ T cells to be significantly enriched in clusters 1, 4 and 5 (**Figure 4D**). These clusters were characterized by higher relative expression of proteins linked to glycolysis and lactic acid metabolism such as the glucose transporter GLUT1, glyceraldehyde-3-phosphate dehydrogenase (GAPDH), lactate dehydrogenase (LDH) and monocarboxylate transporter 1 (MCT1) (**Figure 4F**)^55, 56^. Strikingly, mass cytometry again recapitulated the distinct age-dependent gradient of CD38 observed via flow cytometry and RNA-seq, with CB-dominated clusters showing high CD38 expression (**Figure 4G, S4B**).

Compared to CB naïve CD4^+^ T cells, AB cells appeared more reliant on FAO for OXPHOS, as indicated by higher levels of carnitine palmitoyltransferase (CPT1a), an essential enzyme of FAO^57^, and lower levels of proteins associated with glycolysis (**Figure 4F, 4H**). Additionally, adult naïve CD4^+^ T cells exhibited a generally more homogenous metabolic phenotype with the majority falling into the CPT1a-high cluster 2 (**Figure 4C-D**). Although differential expression of proteins associated with glycolysis, lactic acid metabolism, FAO and OXPHOS drove the major age-related differences in metabolic phenotypes, this data also hinted at less pronounced differences in other metabolic pathways. For example, naïve CD4^+^ T cells from CB had higher expression of CD98, which is known to play a role in both T cell development and activation^58^. Meanwhile, AB naïve CD4^+^ T cells had higher levels of the glutamine metabolism enzyme glutamate dehydrogenase (GLUD1), which is of interest given the key role glutamine metabolism plays in a range of T cells functions, including activation (**Figure 4F, 4H**)^58, 59^. Together, these data further indicate that naïve CD4^+^ T cells from CB are metabolically unique compared to their AB counterparts, and that they exhibit key features of glycolytic dependence.

### Neonatal naïve CD4^+^ T cells are more reliant on glycolysis than their adult counterparts

To further test our observation that neonatal versus adult naïve CD4^+^ T cells appear more glycolytically-dependent, we first used SCENITH^60^, a flow cytometry-based method that combines pharmacologic inhibition of metabolic pathways with a puromycin antibody-based readout of cellular ATP production (**Figure 5A**). SCENITH can measure both the pathway by which cells generate energy (i.e. glycolysis versus mitochondrial OXPHOS) as well as their preferred energy source (i.e. glucose versus fatty acid and amino acid oxidation). As metabolic state can be impacted by many exogenous factors, we first performed SCENITH on CB and AB CD4^+^ T cells prior to and after cryopreservation. This did not reveal significant differences in relative OXPHOS or glycolysis dependence between naïve CD4^+^ T cells that were processed immediately after collection versus those initially frozen, then thawed and rested overnight prior to analysis (**Figure S5A**).

**Figure 5:**
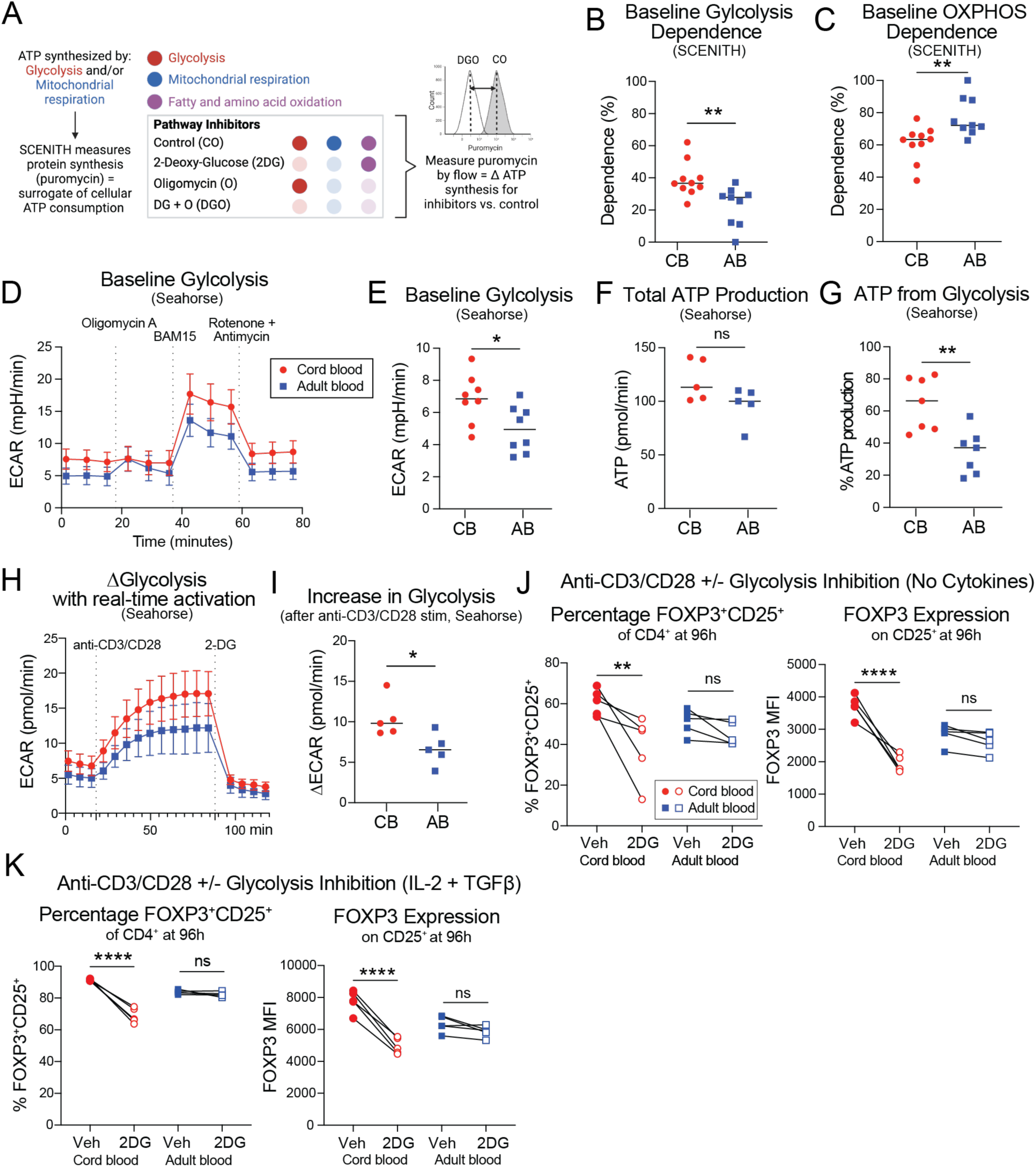
Neonatal naïve CD4^+^ T cells demonstrate a metabolic preference for glycolysis. (A) Schematic of SCENITH workflow. (B,C) Baseline (non-activated) dependence on glycolysis and mitochondrial oxidative phosphorylation in cord blood (CB) and adult blood (AB) naïve CD4^+^ T cells measured via SCENITH. (D) Seahorse graph of ECAR (glycolysis) of baseline (non-activated) CB and AB naive CD4^+^ T cells. (E) Quantification of baseline glycolysis between CB and AB naive CD4^+^ T cells via Seahorse. (F) Total ATP production and (G) percent of ATP made from glycolysis for CB versus AB naive CD4^+^ T cells via Seahorse. (H,I) The change in ECAR for CB and AB naive CD4^+^ T cells after acute TCR activation via real-time introduction of anti-CD3/CD28 soluble antibodies in Seahorse plate reader, between via Seahorse. (J,K) Quantification of the percentage change in FOXP3^+^CD25^+^ and FOXP3 MFI of CB and AB cells after 96hr ImmunoCult^TM^ stimulation of naïve CD4^+^ T under no cytokine or IL-2+TGF-β conditions with or without concurrent inhibition of glycolysis via treatment with 0.5mM 2DG. Vehicle (Veh) control = DMSO. All data points represent individual samples from different biological donors. B, C, and E are two combined experimental replicates. D, F-K are one representative experiment of two experimental repeats.

Based on these results, we opted to perform our metabolism-focused experiments on cryopreserved samples to enable parallel processing and analysis of multiple donors. First, we used SCENITH to compare naïve CD4^+^ T cells from CB or AB at baseline, prior to activation. Mean puromycin fluorescence intensity was equivalent between groups in the absence of metabolic inhibition, suggesting relatively equivalent ATP use irrespective of age (**Figure S5B-C**). Changes to puromycin incorporation in the setting of various pathway inhibitors revealed CB naïve CD4^+^ T cells to have an increased relative dependence on glycolysis versus AB, with a converse decrease in relative dependence on OXPHOS (**Figure 5B-C**). We did not observe any differences in energy substrate preference by age, with both CB and AB naïve CD4^+^ T cells demonstrating a higher reliance on fatty acid and amino acid oxidation versus glucose, as is expected for non-activated, resting cells (**Figure S5D**).

To validate our results from SCENITH and further measure the absolute level of energy pathway use between CB versus AB, we performed Seahorse on resting naïve CD4^+^ T cells. This showed higher baseline extracellular acidification rate (ECAR) in CB cells, reflective of increased glycolysis (**Figure 5D-E**). Consistent with puromycin levels measured by SCENITH, Seahorse did not show an age-dependent difference in total ATP generation by naïve CD4^+^ T cells, but a relative increase in ATP made from glycolysis in CB was confirmed (**Figure 5F-G**).

Seahorse did not reveal differences in baseline oxygen consumption rate (OCR, measurement of OXPHOS), ATP from OXPHOS, or spare respiratory capacity between ages (**Figure S5E-H**). Taking advantage of the Seahorse Human T cell Activation Assay, we also were able to measure ECAR and OCR continuously in the two hours following anti-CD3/CD28 activation with Immunocult^TM^. This showed that CB naïve CD4^+^ T cells increase their glycolytic activity even more than AB following TCR stimulation, which was matched in this setting by age-specific increases in OXPHOS (**Figure 3H-I, S3I-J**).

Considering the integral relationship between T cell metabolism and function, we next probed whether the glycolytic state of CB naïve CD4^+^ T cells was linked to their observed heightened capacity for FOXP3 expression upon stimulation. For this, we employed 2-deoxy-D-Glucose (2DG), a pharmacological inhibitor of glycolysis, which we titrated to a dose of 0.5mM that did not impact CD4^+^ cell viability, proliferative capacity (Ki67), or activation (CD69) during 96 hour stimulation (**Figure S5K-M**). We then treated CB and AB naïve CD4^+^ T cells with 2DG in tandem with “No Cytokine” or “iTreg” culture conditions for 96 hours. 2DG treatment led to a significant reduction in the percentage of FOXP3^+^CD25^+^ cells generated from CB naïve CD4^+^ T cells, as well as their level of FOXP3 expression, under either culture condition, without analogous effects on AB (**Figure 5J-K**). Taken together, this data confirms a higher baseline glycolytic dependency among CB versus AB naïve CD4^+^ T cells and connects this with their heightened ability to upregulate FOXP3 upon stimulation.

### CD38 inhibition limits glycolysis and FOXP3 expression by CB naïve CD4^+^ T cells

To dissect if CD38 expression on CB versus AB naïve CD4^+^ T cells was functionally linked with their heightened glycolytic dependency and capacity to upregulate FOXP3, we took advantage of the potent and specific CD38 pharmacological inhibitor, 78c^20, 23^. First, we incubated CB and AB naïve CD4^+^ T cells with 10uM 78c for 24 hours prior to assessment of their metabolic state via Seahorse. CD38 inhibition led to a significant reduction in glycolysis by CB naïve CD4^+^ T cells, as measured by ECAR, a decrease in ATP from glycolysis (**Figure 6A-B**), and an accompanying increase in their OXPHOS activity (**Figure S6A**). Analogous treatment of AB naïve CD4^+^ T cells with 78c did not lead to metabolic changes in any of these metabolic parameters (**Figure S6B**). Additional titration of the 78c concentration followed by measurement of glycolysis via SCENITH confirmed a dose-dependent decrease in CB glycolysis dependency without effects on AB (**Figure S6C**).

**Figure 6:**
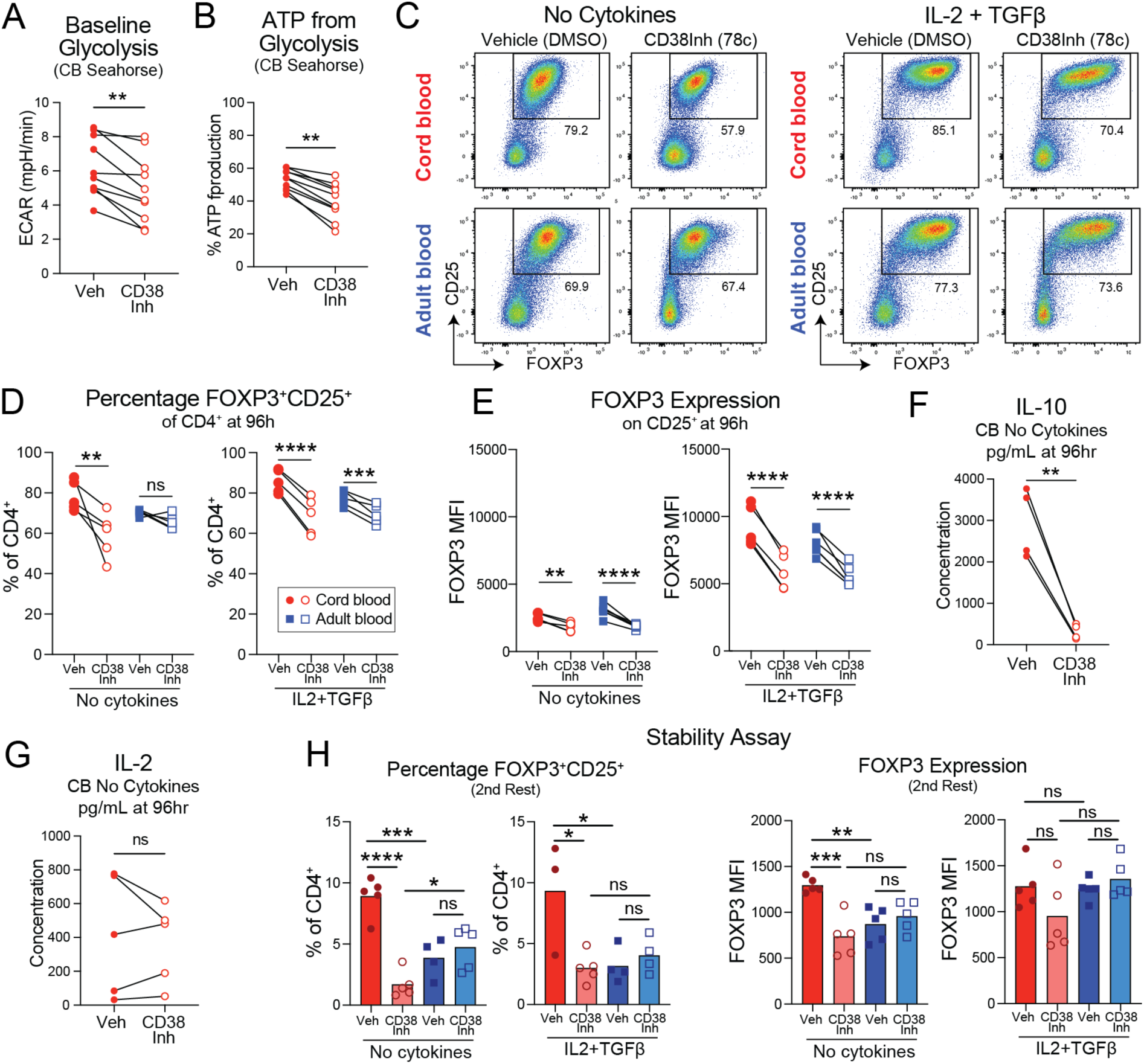
Inhibition of CD38 activity in cord blood naïve CD4^+^ T cells reduces their FOXP3 expression and glycolytic capacity. (A) Quantification of ECAR as a measure of baseline glycolysis between vehicle (DMSO) and CD38 inhibitor (5uM 78c) treated CB naive CD4^+^ T cells via Seahorse. (B) Percent of ATP made from glycolysis for DMSO versus 78c treated CB naive CD4^+^ T cells via Seahorse. (C-E) Representative flow plots and graphs of the percentage of FOXP3^+^CD25^+^ and FOXP3 MFI on FOXP^+^CD25^+^ cells from cord blood (CB) and adult blood (AB) 96hr stimulated cells with vehicle (DMSO) or CD38 inhibitor (5uM 78c) under “No Cytokine” or “IL-2+TGF-β” conditions. Cells were treated with DMSO or 78c for 24hr prior to activation and measured at 96hr after stimulation. (F,G) ELISA- based measurement of cytokines in cell culture supernatant of DMSO or 78c treated CB stimulated naïve CD4^+^ T cells at 96hr from “No Cytokine” condition. (H) FOXP3^+^CD25^+^ stability assay under “No Cytokine” or “IL-2+TGF-β” conditions where 78c was added for 24hr prior to activation and during the first 96hr of activation but was then washed out and not re-added for the duration of the stability assay. All data points represent individual samples from different biological donors. A-E display one representative experiment from two or three repeats. F-H display one experiment.

We then extended treatment with 78c through the full 96 hours of stimulation with or without IL-2 and TGFβ. CD38 inhibition under “No Cytokine” conditions resulted in a significant decrease in the percentage of FOXP3^+^CD25^+^ cells from CB but not AB. In 78c-treated “iTreg” cultures there was a decrease in the percentage of FOXP3^+^CD25^+^ cells in both age groups. The intensity of FOXP3 expression among FOXP3^+^CD25^+^ cells was blunted by CD38 inhibition irrespective of age or culture condition (**Figure 6C-E**). Although 78c did not limit CD69 expression on AB or CB CD4^+^ T cells at the concentration used (**Figure S6D**), we did observe reduced Ki67 expression at 96 hours as marker of recent cell proliferation across all groups (**Figure S6E**), an effect that was not observed among FOXP3^+^CD25^+^ cells specifically (**Figure S6F**). Further titration of the 78c concentration under “No Cytokine” conditions showed no effect on cell viability. There was differential sensitivity to 78c between CB and AB with regard to percentage of FOXP3^+^CD25^+^ cells and their FOXP3 expression. Whereas a significant reduction for each parameter was seen in CB at 5uM 78c, similar effects were achieved in AB only at 40uM. In contrast, there was an equivalent dose-dependent effect irrespective of age on Ki67, suggesting that 78c-mediated inhibition of FOXP3 expression in CB was not merely a consequence of inhibited proliferation (**Figure S6D**).

To further explore the functional impact of CD38 inhibition on the tolerogenic potential of CB naïve CD4^+^ T cells, we measured cytokine production in 96 hour supernatant from “No Cytokine” cultures. This revealed a decrease in IL-10 but not IL-2 in 78c-treated CB (**Figure 6F-G**). We additionally performed FOXP3 stability assays as previously detailed, this time comparing AB and CB cells that were initially cultured with or without 78c in either “No Cytokine” or “iTreg” conditions. Again, there were no statistically significant differences seen at the “first rest” timepoint (**Figure S6H**). However, at the “second rest” timepoint, we observed significant reductions in both the percentage of FOXP3^+^CD25^+^ cells and the FOXP3 MFI in 78c-treated “No Cytokine” CB cultures. Under “iTreg” conditions, CD38 inhibition likewise led to reduced CB FOXP3^+^CD25^+^ cells, with a non-significant downward trend seen in their FOXP3 expression. 78c had no effects on either parameter among AB cells, irrespective of culture condition (**Figure 6H**).

Based on these results, we wondered if CD38 inhibition would lead to preferential effects on CB versus AB irrespective of differentiation trajectory. Our bulk RNA-seq data revealed higher transcript levels for type 2 cytokines, i.e. IL-4, IL-5, IL-13, in stimulated CB naïve CD4^+^ T cells. Based on this observation and prior literature suggesting a role for glycolysis in promoting Th2 differentiation^61, 62^, we thus opted to examinate the relative effect of 78c on CB versus AB under Th2 differentiation conditions. CB and AB naïve CD4^+^ T cells were stimulated via plate coated anti-CD3/CD28 antibodies and treated with IL-4 and anti-IFNy. After 3 days, TCR stimulation was removed and cells were rested overnight with IL-2, IL-4, and anti-IFNy before with PMA/Ionomycin treatment and flow cytometry analysis. These conditions generated IL-4^+^IL-13^+^ cells from both CB and AB, with an equivalent percentage of double-producers seen across both ages (**Figure S6I**). In contrast to the preferential effect seen on FOXP3 expression in CB with CD38 inhibition, treatment with 78c for 24 hours prior to and during Th2 differentiation led to an equivalent reduction in the percentage of IL-4^+^IL-13^+^ cells in both CB and AB. Thus, while CD38 inhibition can likewise impact naïve CD4^+^ differentiation down a Th2 trajectory, heightened CB sensitivity to CD38 inhibition is not seen in this setting as it is for FOXP3 expression following TCR stimulation.

### The CD38-NAD-SIRT1 axis modulates FOXP3 expression in stimulated neonatal naïve CD4^+^ T cells

We next sought to further dissect how CD38 might functionally shape the metabolic state and Treg potential of CB naïve CD4^+^ T cells. CD38’s enzymatic function in catalyzing the synthesis of ADP ribose and cADPR has been shown to deplete cellular NAD+ stores^29^. Consistent with this being a relevant mechanism in our studies, we observed lower NAD+ levels in naïve CD4^+^ T cells from CB as compared to AB (**Figure 7A**). Furthermore, pharmacological inhibition of CD38 in CB naïve CD4^+^ T cells led to an increase in NAD+ levels **(Figure 7B**). NAD+ can impact cellular function through multiple avenues, including its role as a cofactor for oxidoreductases, an electron donor for redox reactions and a substrate for the sirtuin family of deacetylases ^63^. We were particularly intrigued by this third possibility, given existing literature linking the sirtuin family member, SIRT1, to T cell metabolism and FOXP3 expression. Specifically, low SIRT in terminally differentiated human memory CD8^+^ T cells was shown to facilitate their glycolytic state^64^. Separately, SIRT1-mediated histone deacetylation has been shown to decrease chromatin accessibility of the *FOXP3* locus^65^, and SIRT1 deacetylation of FOXP3^32, 33^ can target the protein for degradation. To further probe the relevance of histone-mediated effects, we assessed global levels of H3K27ac in CB versus AB naïve CD4^+^ T cells, a histone modification associated with heightened *FOXP3* induction^66^. Antibody staining, however, did not reveal any age-related differences in H3K27ac levels (**Figure S7A**). We then performed a flow-based proximity ligation assay to measure acetylation of FOXP3 protein. This revealed significantly higher levels of acetylated FOXP3 protein in CB versus AB naïve CD4^+^ T cells (**Figure 7D**). This result is consistent with a model in which lower SIRT1 activity in CB leads to less FOXP3 deacetylation and, thus, higher FOXP3 levels due to its reduced degradation.

**Figure 7:**
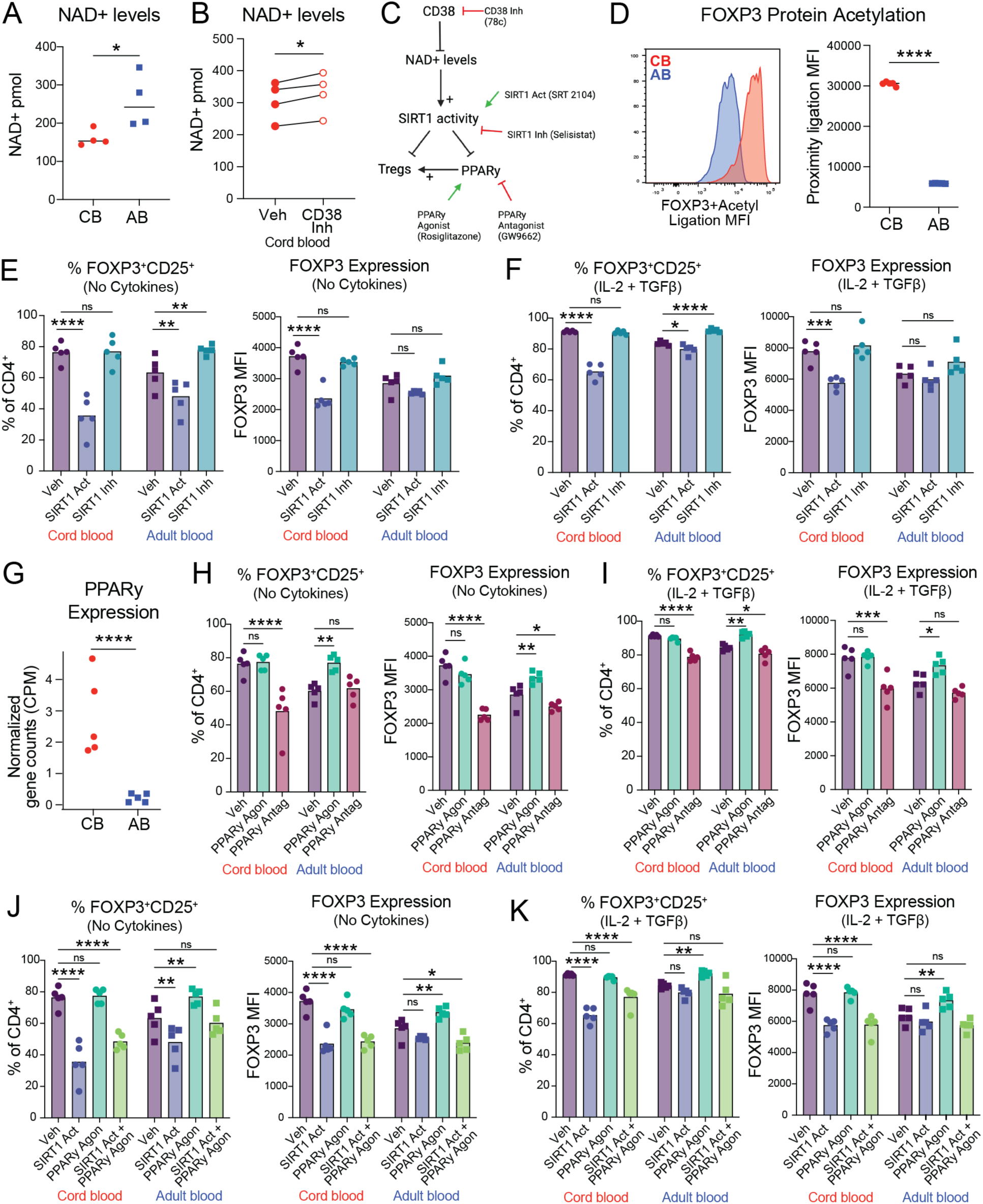
The NAD+ dependent deacetylase, SIRT1, modulates FOXP3 expression in stimulated neonatal naïve CD4^+^ T cells. (A) NAD+ levels measured between cord blood (CB) and adult blood (AB) cell lysates from naïve CD4^+^ T cells at baseline. (B) NAD+ levels measured between DMSO or 78c 24hr treated CB cell lysates from naïve CD4^+^ T cells at baseline. (C) Schematic of CD38-NAD-SIRT1-Tregs-PPARy pathway with pharmacological perturbations used in subsequent experiments included. (D) Acetylation of FOXP3 protein in non-activated naïve CD4+ T cells from CB and AB was measured via a proximity ligand assay (PLA) where anti-FOXP3 and anti-acetylated lysine antibodies were added and their proximity to each other measured via flow cytometry to assess acetylation levels on FOXP3 protein by age. (E,F) CB and AB naïve CD4^+^ T cells were treated with vehicle control (DMSO), SIRT1 activator (20uM of SRT 2104), or SIRT1 inhibitor (20uM of Selisistat) for 24 hours prior to activation under no cytokine or IL-2+TGF-β conditions and FOXP3^+^CD25^+^ and FOXP3 MFI was measured 96hr after activation. (G) Normalized gene counts of PPARy (PPARG gene) at 48hr after activation from bulkRNAseq data. (F) CB and AB naïve CD4^+^ T cells were treated with vehicle control (DMSO) or PPARy agonist (60uM of Rosiglitazone) for 24 hours prior to activation and Treg measurement 96hr after activation. (H,I) CB and AB naïve CD4^+^ T cells were treated with vehicle control (DMSO) or PPARy antagonist (2.5uM of GW9662) for 24 hours prior to activation under no cytokine or IL-2+TGF-β conditions and FOXP3^+^CD25^+^ and FOXP3 MFI was measured 96hr after activation. (J,K) CB and AB naïve CD4^+^ T cells were treated with vehicle control (DMSO), SIRT1 activator (20uM of SRT 2104), PPARy agonist (60uM of Rosiglitazone), or both SIRT1 activator and PPARy agonist for 24 hours prior to activation under no cytokine or IL-2+TGF-β conditions and FOXP3^+^CD25^+^ and FOXP3 MFI was measured 96hr after activation. All data points represent individual samples from different biological donors. A, B, D, E, F, H-K show one experimental replicate of two.

To test the relevance of SIRT1 in FOXP3 expression in stimulated CB naïve CD4+ T cells, we treated CB and AB naïve CD4^+^ T cells with a SIRT1 Activator (SRT 2104), a SIRT1 Inhibitor (Selisistat), or DMSO as a vehicle control for 24 hours prior to activation in “No Cytokine” or “iTreg” conditions. After 96 hours, SIRT1 activation under either culture condition led to significantly reduced percentages of FOXP3^+^CD25^+^ cells in CB as well as their lower expression of FOXP3. SIRT1 activation led to more modest but still statistically significant reductions in the percent of AB FOXP3^+^CD25^+^ cells, without significant impact on their FOXP3 MFI. In contrast, SIRT1 inhibition did not impact FOXP3 expression in CB, but in AB did lead to modest but statistically significant increases in the percent of FOXP3^+^CD25^+^ cells (**Figure 7E-F, S7B**). This supports the idea that very low SIRT1 activity in CB naïve CD4^+^ T cells contributes to their increased FOXP3 expression, whereas AB cells have at baseline somewhat higher SIRT1 activity, which can be further modulated pharmacologically to tune their tolerogenic potential. In other cell types, SIRT1 is known to engage in a negative feedback loop with the nuclear hormone receptor PPARγ^67, 68^. PPARγ is a pleiotropic transcription factor best known for its role in adipogenesis, lipid metabolism and insulin sensitivity^69^. It has also been shown to have a key role in the biology of fat-resident Tregs^70^ as well as in Treg generation and function more broadly^71–73^. In our RNA-seq dataset, PPARγ gene counts were about 10-fold higher in 48 hour stimulated CD4^+^ T cells from CB versus AB (**Figure 7G**). To test if increased neonatal PPARγ expression could also contribute to increased early life Treg generation, we treated naïve CD4^+^ T cells with the PPARγ agonist Rosiglitazone, the PPARγ antagonist GW9662, or DMSO vehicle control for 24 hours prior to activation in “No Cytokine” or “iTreg” conditions. Measurement of CB FOXP3^+^CD25^+^ percent of FOXP3 MFI at 96 hours revealed no change with PPARγ agonism in either culture condition, but PPARγ antagonism did lead to a significant reduction of both parameters in either culture condition. Conversely, PPARγ agonism with Rosiglitazone in AB significantly increased the percent of FOXP3^+^CD25^+^ cells as well as their FOXP3 expression level to proportions more commensurate with CB. Conversely PPARγ inhibition of AB via GW9662 decreased FOXP3^+^CD25^+^ cells as well as their FOXP3 expression (**Figure 7H-I**). These data suggest that differential age-dependent PPARγ expression in naïve CD4^+^ T cells may contribute to their distinct capacities for FOXP3 expression.

Finally, we sought to clarify if CD38 levels and, by extension, SIRT1-modulation of FOXP3 expression in naïve CD4^+^ T cells was mediated primarily through regulation of PPARγ. To dissect this, we pre-treated naïve CD4^+^ T cells prior to activation with our SIRT1 activator, PPARγ agonist, or both. Interestingly, while we recapitulated reduced CB and AB FOXP3^+^CD25^+^ percentages and lower CB FOXP3 MFI with SIRT1 activation alone, addition of Rosiglitazone to agonize PPARγ in this setting only led to only a very modest restoration of CB FOXP3^+^CD25^+^ percentages and was insufficient to increase CB FOXP3 expression levels (**Figure 7H**). Of note, titration of the pharmacological agents used for these experiments did not reveal substantial effects on CB or AB cell viability, activation or proliferation, with the exception of a small dose-dependent decrease in Ki67 expression in the setting of SIRT1 agonism. Likewise, effects of SIRT1 agonism and PPARγ in reducing CB FOXP3^+^CD25^+^ percentages were also achieved at even lower concentrations of each agent (**Figure S7G-R**). Collectively, these results indicate that the CD38-NAD-SIRT1 axis works through multiple different mechanisms to influence the CD4^+^ T cell-intrinsic capacity for early life FOXP3 expression. While PPARγ may be one downstream pathway, it is likely that direct effects of SIRT1 on FOXP3 protein stability, and possibly the metabolic state of naïve CD4^+^ T cells are also critical. Together, these studies provide significant new mechanistic insights into how heightened neonatal expression of CD38 in the naïve CD4^+^ T cell compartment helps enable their regulatory potential and, by extension, early life window for tolerance.

## Discussion

Here we demonstrate that early life naïve CD4^+^ T cells have a unique metabolic state, characterized by a higher reliance on glycolysis, and that this is mediated at least in part by their high expression of CD38. In contrast, the adult naïve CD4^+^ compartment is characterized by negligible CD38 expression, which corresponds to lower glycolytic dependence. We further show that naïve CD4^+^ T cells from CB versus AB display a heighted capacity to upregulate FOXP3 upon stimulation and that, specifically under iTreg conditions, they display greater suppressive capacity, demethylation of CNS2, and stability of FOXP3 expression. Inhibiting glycolysis or CD38 during stimulation preferentially decreased FOXP3 expression in stimulated CB versus AB naïve CD4^+^ T cells, implicating CD38 as a key age-dependent regulator. Through further pharmacological inhibition, we demonstrate that CD38’s role as an NADase in neonatal CD4^+^ T cells potentially limits SIRT1 deacetylation, which in turn could contribute to increased tolerogenic potential, likely via several mechanisms including modulation of PPARγ expression, cell metabolic state, as well as direct tuning of FOXP3 protein expression. Collectively, this work elucidates a central paradigm governing neonatal T cell development of regulatory properties, which may functionally enable the early life window of opportunity for establishing peripheral immune tolerance.

Measuring Treg generation from human naïve T CD4^+^ T cells is fraught with the challenge that FOXP3, CD25, CTLA4 and other Treg “markers” are also expressed by activated effector T cells. “iTreg” culture conditions with IL-2 and TGF-β, are currently the gold standard for generating human Tregs ex vivo. However, these iTregs perform sub-optimally in assays of Treg function and do not demonstrate as extensive CNS2 demethylation or FOXP3 protein stability as compared to endogenous Tregs from human blood or tissue^74^. This has led to consideration of alternative approaches in pursuit of tolerogenic cellular therapy, such as endogenous or exogenous Treg expansion, in cellular therapy. Just recently, however, a CRISPR screen identified the RBPJ–NCOR complex as a targetable repressor of FOXP3 stability in iTregs^49^. In this context, our observation that iTreg conditions enable CB naïve CD4^+^ T cells to develop heightened suppressive capacity, CNS2 demethylation, and FOXP3 stability when compared their adult counterparts is particularly intriguing.

These observations open new questions about what regulates CD38 expression on naïve CD4^+^ T cells, and how it shifts in an age-dependent manner. PAXIP1 (also referred to as Pax-interacting protein 1 or PTIP) has been shown in macrophages to promote CD38 expression by cooperatively increasing H3K27 acetylation of an intronic enhancer^75^. PAXIP1 has not been specifically studied as a regulator of CD38 in T cells but does have established roles in the early stages of T cell development, specifically enabling genomic stability during TCR gene re-arrangement and promoting expression of S1PR1 on single-positive thymocytes to enable their thymic egress^76^. Whereas PAXIP1 is not known to have differential expression by age, the RNA-binding protein, Lin28b, is specifically increased in early life hematopoietic progenitor cells, where it is thought to shape many features of fetal and neonatal lymphocytes^19, 77^. Studies in separate cell types have also shown that Lin28 promotes glucose metabolism via its inhibition of the let-7 microRNA and downstream activation of MTOR signaling^78^, making it another intriguing candidate for shaping age-dependent CD38 expression.

Work showing that CD38 is enriched on recent thymic emigrants (RTEs)^25^ provides an intriguing explanation for why CD38 expression might be increased in the early life window, a period demarcated by high thymic output^79^. However, rather than bimodal expression of CD38 in the neonatal naïve CD4^+^ compartment, we observe that the mean CD38 expression shifts down with age for the entire population. A slow decrease in CD38 levels among circulating CD4^+^ T cells following thymic egress, in tandem with diminishing thymic output could lead to this pattern of expression. Additionally, although we found CD38 to be elevated in CD31^+^ versus CD31^neg^ cells in both CB and AB, large age-dependent differences were preserved in both CD31^+^ and CD31^neg^ cells, suggesting that heightened early life CD38 expression is not merely a reflection of RTE frequency. Thus, further studies are needed to define key regulators of CD38 expression in T cells and how this relates to the age at which they were generated and the length of time they have spent in circulation.

Beyond CD38 specifically, our work underscores the concept that the metabolic state of early life lymphocytes fundamentally shapes their functional capacity^39, 80^. We show that human naïve neonatal CD4^+^ T cells exhibit higher relative dependence on glycolysis than their adult counterparts, analogous to findings in human CD8^+^ T cells^81^. Prior work in murine models has linked the glycolytic state of neonatal CD8^+^ T cells to heightened effector memory differentiation and a reduced capacity for generation of central memory subsets^19^. Here we show that reducing the glycolytic state of CB naïve CD4^+^ T cells, either directly via 2DG or indirectly via 78c, diminishes their capacity for Treg generation. Further work is needed, however, to decipher the mechanistic basis of this link between naïve CD4^+^ T cell metabolism and function. One possibility based on the literature^56^ is that a high glycolytic state reduces activity of myc-promoted binding protein (MBP1), a transcription factor known to repress of a *FOXP3-*E2 splice variant^82^. Glycolysis-facilitated release of this repression may increase expression of this splice variant previously shown to promote Treg differentiation and stability^83^.

While there is a given the body of work illustrating the importance of OXPHOS in the function of fully differentiated Tregs ^84, 85^, this literature is generally surrounding terminally differentiated Tregs from adult donors. Not only have some publications shown glycolysis to be critical for iTreg induction and suppressive function^82^, but there are minimal studies on the metabolic requirements for iTreg and Treg induction from early life or neonatal T cells. We found that early life naïve CD4^+^ T cells are preferentially reliant on glycolysis compared to adult cells, and that this glycolytic preference translates into functional outcomes (i.e. glycolysis inhibition reduces the percentage of FOXP3^+^CD25^+^ cells generated from stimulation of neonatal but not adult naïve CD4^+^ T cells). Additionally, CD38 is known to negatively impact mitochondrial oxidative phosphorylation^23^. Thus, high CD38 expression in CB naïve CD4^+^ T cells may be part of this CD38-glycolysis/oxidative phosphorylation axis, thus providing an age dependent unique mechanism by which tolerance is promoted in early life. We find our study uniquely positioned to understand both the age-dependent differences in T cell metabolic requirements, as well as how those differences impact emerging cells, rather than terminally differentiated ones. However, additional work will be needed to disentangle the longer-term fate of Tregs emerging from the neonatal naïve CD4^+^ T cell compartment. Examination of environmental impacts, such as the impact of changing oxygen tension at birth, as well as the development of animal models, analogous to those used to study neonatal versus adult CD8^+^ T cells^46^, will hopefully help determine if the glycolytic tone accompanying the induction of Tregs from neonatal CD4^+^ T cells negatively impacts their stability of function.

Through pharmacological inhibition and activation of SIRT1, we identified this NAD-dependent deacetylase as a potential key downstream mediator of CD38’s influence on FOXP3 expression in neonatal naïve CD4^+^ T cells. While SIRT1 activity is known to increase in aging cells and mediate physiological changes associated with healthy aging, most literature has focused on differential activity between mid and late adult life, with little attention paid thus far to specific repression of its activity in the neonatal window^86^. Given the wide range of histone and non-histone SIRT1 acetylation targets, changes in multiple cell pathways likely contribute to SIRT modulation of T cell behavior. Thus, disentangling their relative impact in neonatal T cells will take extensive work. Based on our data showing differential FOXP3 protein acetylation in CB versus AB, SIRT1-mediated FOXP3 deacetylation^87^ is one candidate mechanism by which CD38 may influence FOXP3 expression in CB that is deserving of further study. However, SIRT1 may also influence early life T cell behavior by augmenting glycolysis over OXPHOS via key intermediary transcription factors, e.g. PGC-1α, HIF-1α, or enzymes, e.g. PGAM1^88–90^. We also show a potential role for PPARγ as another SIRT1 interactor that could influence both CD4^+^ T cell metabolism and Treg potential^73, 91^. SIRT1 has been shown to inactivate PPARγ via its deacetylation^67^. Thus, low SIRT1 activity in neonatal naïve CD4^+^ T cells might contribute to higher PPARγ activity and Treg-promoting function. Additionally, PPARγ can inhibit SIRT1 expression at the transcriptional level providing a potential negative feedback loop that could serve as another modulator of SIRT1 activity neonatally in addition to low NAD+ levels^68^.

In conclusion, our findings highlight the translational potential of modulating CD38 expression and glycolytic state in neonatal naïve CD4^+^ T cells to influence their FOXP3 expression. By characterizing CD38 as a metabolic regulator within neonatal T cells, we reveal a potential target for therapeutic strategies aimed at enhancing peripheral tolerance in early life, which could be especially relevant for preventing immune disorders such as autoimmunity and allergy. Furthermore, our work underscores the need for age-specific approaches in immune modulation, as metabolic interventions to enhance Treg induction in neonates may differ substantially from those required in adults. The unique neonatal environment, with its distinct metabolic cues and regulatory proteins like CD38, provides an essential opportunity to understand and leverage the age-dependent plasticity of immune cells. Translationally, future studies targeting these pathways could refine interventions in neonatal immune regulation, advancing both preventive and therapeutic strategies for immune tolerance in early life.

## Limitations of the study

Due to constraints in human sample collection, obtaining sufficient quantities of post-natal infant blood proved challenging. Consequently, cord blood was utilized in this study as a surrogate for early life samples. Although we were able to confirm the persistently high expression of CD38 in a limited number of pre- and post-natal samples, we were unable to fully validate all findings in these samples. This limitation, coupled with the scarcity of post-natal samples, restricted our capacity to examine graded changes in CD38 expression, metabolic state, and Treg potential within the post-natal period to assess temporal dynamics. Collecting early life (0-18 years) human samples is an ongoing collaborative effort and will form the basis of future studies focused on delineating the timing and progression of these age-related observations.

In this study, we used pharmacological agents rather than genetic approaches to dissect the functional role of CD38 in neonatal CD4^+^ T cells. This choice was informed by observations that CRISPR electroporation of cord blood naïve CD4^+^ T cells impaired their activation capacity, even when using a scramble control without CD38 disruption. Moreover, genetic disruption via CRISPR requires cell activation, which would not allow us to assess age-dependent differences in a resting state. Therefore, we opted to use pharmacological compounds that have been well-validated and further optimized the dosing and treatment conditions within our system. Efforts to refine CRISPR techniques for cord blood naïve CD4^+^ T cells are ongoing to enable genetic manipulation of CD38 and its downstream mediators. Finally, future work is needed to determine whether or to what extent CD38 expression influences other aspects of naïve CD4^+^ T cell differentiation. We show here that CD38 inhibition had equal inhibitory effects on Th2 differentiation in both CB and AB, in contrast to the age-specific effects on FOXP3 expression, but we did not fully explore effects on Th1 or Th17 differentiation.

## Supporting information

Supplemental Table 1 - CyTOF Antibody List

## Acknowledgements

We thank Rafael Argüello Ph.D. for contribution of SCENITH reagents, Joanna Halkias M.D., Ph.D., Ari Molofsky M.D., Ph.D., Michael Rosenblum M.D., Ph.D. and Qizhi Tang Ph.D. for helpful discussions. Umbilical cord blood units utilized in this project were obtained from the Umbilical Cord Blood Collection Program, made available under the California Health & Safety Code §§ 1627-1630, and maintained by the Institute for Regenerative Cures, University of California at Davis, Sacramento, CA 95817. Infant blood was obtained from a collaboration with the PROMPT/PREMO Cohort with assistance from NIH Grant R01 HD102381-01 and UCSF’s Benioff Center for Microbiome Medicine. Prenatal spleen samples were obtained in collaboration with the Rutishauser Lab. We acknowledge UCSF’s Parnassus Flow CoLab (RRID:SCR_018206) for assistance in flow cytometry, CyTOF and cell sorting, supported by NIH grants P30 DK063720 and 1S10ODO18040-01. We thank Sandler Program for Breakthrough Biomedical Research Technologies, Methodologies & Cores and UCSF NORC grant (P30DK098722) for funding and use of the Seahorse machine for metabolism measurements. L.D. was supported by funds from the UCSF Immunology T32 Training Grant (NIH grant T32AI007334-33), an NIH NRSA fellowship (1F31AI186359-01A1), and a Trainee Research Award from UCSF’s Benioff Center for Microbiome Medicine. Work was supported by startup funds from UCSF Department of Dermatology and a CoProject grant from the UCSF Immunox Initiative.

## Author Contributions

L.D. and T.C.S. designed the studies and wrote the manuscript. L.D. performed the experiments and analyzed the data. A.M.D performed all Mass Cytometry experiments and analysis. S.C. performed analysis of bulk RNA-seq data. V.G. assisted in experimental design and execution. C.C. performed COMPASS analyses. E.E.R., S.P.O, and L.L.J.P. provided access to preterm birth samples. S.V.L., R.L.R., A.W. and A.J.C provided resources and input on experimental design. T.C.S. oversaw all study design and data analysis. All authors discussed results and commented on the manuscript.

## STAR METHODS

### Experimental Model and Subject Details

#### Human Samples

Deidentified human cord blood samples were purchased University of California, Davis from the California Umbilical Cord Blood Collection Program (UCBCP). Donors were screened for healthy full-term birth, normal weight and no commodities in mother or infant. Healthy non-obese, non-smoking adult donors of ages 18-45 were consented for blood donation under IRB Number 12-09489 and IRB Number 13-11307.

Preterm infant blood samples were obtained from UCSF participants in the Prediction of Maturity, Morbidity, in Preterm Infants (PROMPT) study. This is a multi-site NIH study funded by the NIH (R01 HD102381-01) with a common IRB (together with the University of Iowa) approved by the Western Institutional Review Board (WCG IRB), IRB number 20202748, and conducted in accordance with all applicable ethical guideline and regulations for human subject research. Infant peripheral blood samples were taken from 4 preterm infants born at 30 weeks, with the peripheral blood samples were taken at 6 weeks post-birth.

Fetal spleen samples from 21-24 gestational week-old donors were acquired through the Women’s Options Center at Zuckerberg San Francisco General Hospital after elective termination of pregnancy. Written informed consent was obtained with IRB-approval from, and under the guidelines of the UCSF Human Research Protection Program. Samples were excluded in the cases of (a) known maternal infection, (b) known intrauterine fetal demise, and/or (c) known or suspected chromosomal abnormality. Samples were fully de-identified, had no associated Personal Heath Information (PHI), and researchers had no access to PHI.

Cord blood samples were shipped overnight, and adult and infant blood samples were obtained, left on a rocker at room temperature, and processed within 12 hours of collection. All samples were then uniformly processed using a Ficoll-Plaque gradient to isolate peripheral blood mononuclear cells (PBMCs). PBMCs were frozen at a concentration of 10mil/1mL in freezing media (90% FBS, 10% DMSO) and were then stored in a liquid nitrogen tank. Fetal spleen were digested with media supplemented with 3mg/ml Collagenase IV and 1mg/ml DNase I at 37c for 30-45min in a shaking water bath set to 175rpm. After digestion, they were filtered and pelleted, after which the lymphocyte layer was isolated via Ficoll gradients. PBMCs were frozen, stored, and re-thawed similarly to CB. Sample collection was attentive in considering sex as a biological variable and included equal proportions of cord blood and adult blood samples from male and female donors.

### Method Details

#### Cell culture and stimulation

Cells were thawed and washed before resting overnight in a 37C 5% CO2 incubator in Resting Media (RPMI with 10% FBS, 1% 100x Pen/Strep, 1% Hepes, and 1% Glutamax). After resting, naïve CD4^+^ T cells were isolated using the StemCell EasySep™ Human Naïve CD4^+^ T Cell Isolation Kit. For stimulation assays on which RNA sequencing was performed (see details below), live naïve CD4^+^ T cells were isolated by fluorescence-activated cell sorting (FACS) based on the following markers: CD4^+^CD8^-^CD45RA^+^CD27^+^CCR7^+^CD95^-^. Naïve CD4^+^ T cells were either used immediately or activated and cultured with STEMCELL ImmunoCult™ Human CD3/CD28 T Cell Activator in STEMCELL Immunocult^TM^ T cell Media (+ 0.1% Pen/Strep) at a concentration of 100,000 cells/200uL media in 96 well U bottom plates in a 37C 5% CO2 incubator for up to 96 hours post-activation. For “No cytokine” culture conditions, no cytokines were added to the culture. iTreg culture conditions were adapted from prior work^92^, and included addition of exogenous IL-2 (10 ng/ml; PeproTech) and TGF-β (50 ng/ml; PeproTech). 100uL media was removed and replaced with fresh media with no cytokines or IL-2+TGF-β. To verify appropriate activation, ImmunoCult™ Human CD3/CD28 T Cell Activator was compared to other standardized protocols for human T cell activation: Dynabeads™ Human T-Activator CD3/CD28 for T Cell Expansion and Activation, coating the plate in 2ug/mL CD3 and 4ug/mL CD28 antibodies either overnight at 4°C or for 4 hours at 37°C before washing and adding cells for activation.

#### ELISA

Supernatants from cell culture were taken at the time of staining for flow cytometry. 120uL of supernatant for each sample were stored at -80C and sent to Eve Technologies for analysis using their “Human Cytokine Proinflammatory Focused 15-Plex Discovery Assay® Array (HDF15).”

#### SCENITH

SCENITH was performed as described previously^60^. SCENITH reagents (inhibitors, puromycin, anti-puromycin antibodies) were obtained from Rafael Argüello Ph.D. and used according to kit instructions for in vitro human T cell metabolic profiling. Briefly, 100,000 non-activated naïve CD4^+^ T cells or 96 hour activated cells were incubated for 20 minutes with Control (DMSO), 2-Deoxy-Glucose (2DG; 100 mM), Oligomycin (O; 1 µM), or a combination of 2DG and Oligomycin (DGO). Following incubation with inhibitors, Puromycin (final concentration 10 µg/mL) was added to the culture. Due to their lower metabolic state, non-activated cells were incubated with Puromycin for 40 minutes, while 96 hour activated cells were incubated for 15 minutes. After puromycin treatment, cells were washed with cold PBS and stained for flow cytometry in a panel that included an anti-puromycin antibody. Changes in puromycin MFI between conditions were used as a readout for changes in metabolic state when inhibitors of pathways were added.

#### Seahorse

Cells were analyzed using an Agilent Seahorse XFe96 analyzer with the Seahorse XF T Cell Metabolic Profiling Kit. Naïve CD4^+^ T cells were seeded at 150,000 cells/well and kit protocol was followed. Results were analyzed using the Agilent Seahorse Analytics web browser.

#### Flow cytometry

Cells were stained for surface antibodies in PBS + 2% FBS for 30 minutes at 4C. For intracellular staining, cells were fixed and permeabilized using the Foxp3 staining kit (eBioscience, Catalog No. 00-5523-00) buffer for 20 minutes at 4°C then stained in permeabilization buffer for 60 minutes at 4°C. Stained cells were run on a Fortessa (BD Biosciences) in the UCSF Flow Cytometry Core. Flow cytometry data was analyzed using FlowJo software (FlowJo, LLC).

#### Pharmacological perturbations

Chemicals were added after naïve CD4^+^ T cell isolation. Cells were rested for 24 hours with chemicals and were analyzed via Seahorse for non-activated metabolic impacts or were activated with ImmunoCult™ Human CD3/CD28 T Cell Activator after 24 hours with chemicals under “no cytokine” or “iTreg” culture conditions and stained for flow cytometric analysis 96 hours after activation with a 100uL media change at 48HR. The following chemicals were used: 2-Deoxy-Glucose (2DG; Sigma Aldrich, Cat #D3179-250MG), final concentration 0.5mM, CD38 inhibitor 78c (Sigma-Aldrich, Cat #5387630001), final concentration 5uM; SIRT1 inhibitor: Selisistat (SelleckChem, Cat #S1541), final concentration 20uM; SIRT1 activator: SRT 2104 (Medchem Express, Cat #HY-15262), final concentration 20uM; Rosiglitazone (BRL, Cat #49653) a selective PPARγ agonist, final concentration 60uM; GW9662, PPARy antagonist, final concentration 2.5uM.

#### NAD+ level measurement

The NAD/NADH Assay Kit (Colorimetric) (Abcam Cat # ab65348) was used according to the manufacturer’s protocol. Naïve CD4^+^ T cells were isolated from CB and AB PBMCs and lysed. The supernatant from cell lysis was used to perform the assay. Samples were then assessed for total NAD+ on a microplate reader and OD was converted to NAD+/pmol via standard curve. For 78c treatment, naïve CD4^+^ T cells were isolated and treated with 78c or DMSO vehicle control for 24 hours prior to cell lysis and performing the assay.

#### Allogeneic mixed lymphocyte reaction

Naïve CD4^+^ T cells were isolated from CB and AB PBMCs, activated, and cultured under No Cytokine” or “iTreg” culture conditions for 96 hours with a 100uL media change at 48 hours. CD4+CD25+CD127LO and CD4+CD25- cells were then sorted from these cultures along with a thawed and rested overnight universal adult donor’s endogenous CD4+CD25- (poly-MLR) or autologous PBMCs without CD4+CD25+CD127- Tregs (allo-MLR). Responder cells were labeled with CTV and 100,000 responder cells were seeded in 200uL in a 96-well U bottom plate with ImmunoCult anti- CD3/CD28 soluble antibodies (poly-MLR) or 300,000 irradiated PBMCs without CD4+CD25+CD127- from a separate adult donor (allo-MLR). Tregs were titrated into culture at various doses. Proliferation of responder cells was measured at 7 days (allo-MLR).

#### Stability assay

Naïve CD4^+^ T cells were isolated from CB and AB PBMCs, activated with ImmunoCult^TM^ anti-CD3/CD28 antibodies, and cultured under “no cytokine” or “iTreg” culture conditions for 96 hours with a 100uL media change at 48 hours. After 96 hours of activation, cells were washed to remove stimulation and cytokines. Cells were then left to rest for 96 hours with IL-2 added to all conditions and a 100uL media change at 48 hours. After the 96 hour rest, half of the cells were removed and stained for FOXP3 and CD25 expression (“First Rest” samples). The remaining cells were re-stimulated with ImmunoCult^TM^ anti-CD3/CD28 antibodies and IL-2 for 96 hours with a 100uL media change at 48 hours. After 96 hours of re-activation, cells were washed to remove stimulation and cytokines. Cells were then left to re-rest for 96 hours with IL-2 added to all conditions and a 100uL media change at 48 hours. After the second 96 hour rest, the cells and stained for FOXP3 and CD25 expression (“Second Rest” samples). For the stability assay with 78c, 78c or DMSO vehicle control was added for 24 hours prior to the first activation and during the first activation but removed when cells were washed and rested and not added for the remainder of the assay.

#### Th2 differentiation

Naïve CD4+ T cells were isolated from CB or AB PBMCs, seeded (100,000 cells in 200uL media), stimulated via plate coated anti-CD3/CD28 antibodies and treated with IL-4 (12.5ng/mL) and anti-IFNy (10ug/mL). After 3 days after stimulation, 25% of cells were moved to a new plate to remove stimulation and treated with IL-2 (50ug/mL), IL-4, and anti-IFNy. Media was refreshed (100uL) after 24 hours. The following day, cells were stimulated with PMA/Ionomycin for 4 hours with BFA added 2 hours into stimulation. Cells were then stained for flow cytometry to assess cytokine production. For Th2 differentiation with 78c, the same protocol for Th2 differentiation was followed, except naïve CD4+ T cells were treated with the CD38 inhibitor 78c (5uM) or DMSO vehicle control for 24 hours prior to activation and Th2 differentiation.

#### Demethylation

Naïve CD4^+^ T cells were isolated from 5 CB and 5 AB PBMCs, activated, and cultured under “no cytokine” or “iTreg” culture conditions for 96 hours with a 100uL media change at 48 hours. CD4+CD25+CD127LO cells were then sorted from these cultures along with endogenous adult CD4+CD25+CD127LO “Tregs” as methylation control. Samples were shipped as dried pellets to Zymo Research Corporation to undergo targeted bisulfite sequencing of the TSDR loci of the FOXP3 gene. Data was reported as methylation ratios on chromosome coordinates for 15 acetylation sites. Coordinates were mapped to CpG islands within the TSDR loci based on Kressler et al., 2021^93^. A linear mixed effects model was used to analyze the dependence of methylation ratios on the main effects and interactions of CpG site, treatment status (No Cytokine vs. iTreg) and age (CB, AB) using the lme function from the nlme (v3.1.166) package in R (4.4.1(2024-06-14)). A random intercept was used for each patient (∼1 | patient). Type III Analysis of Variance (ANOVA) was performed with Satterthwaite’s method using the anova function from the stats (v4.4.1) package. Post-hoc pairwise comparisons were then performed on the status:age interaction (Pr(>F) = 1.37e-05) condition on CpG site (∼ pairwise ∼status:age | site) using the emmeans package (v1.11.1). P-values at each site were adjusted for family-wise error rate using Tukey’s Honestly Significant Differens (HSD) test.

#### FOXP3 acetylation

The acetylation of FOXP3 protein was assessed using the Duolink^®^ PLA Flow Cytometry Assay (Millipore Sigma, DUO94005) according to manufacturer protocol. Naïve CD4^+^ T cells were isolated from 5 CB and 5 AB PBMCs and treated with antibodies for FOXP3 (Cell Signaling Tech, #12653) and Acetylated-Lysine (Cell Signaling Tech, #9441) at 1:100 dilution for one hour at room temperature. Duolink^®^ PLUS and MINUS PLA probes were then added followed by ligation, amplification, and detection according to manufacturer protocol. Cells were then analyzed by flow cytometry to measure acetylation levels on FOXP3.

#### Statistics

Unless otherwise noted, statistics throughout figures were calculated in Graphpad Prism. For graphs with 2-way comparisons a student’s t-test was used. When CB was compared to AB an unpaired t-test was used and when the same samples (CB or AB) were compared to each other under two different conditions a paired t-test was used – these are denoted by lines linking repeat biological samples. For graphs with three or more comparisons in which biological samples were not repeated a one-way ANOVA with multiple comparison was used. For experiments in which CB or AB from the same donor were subjected to various treatments or measured at different timepoints a repeated measure ANOVA with multiple comparisons was used – these are denoted by lines linking repeated biological samples. Data noted by linked data points reflects paired t-tests, with Bonferroni adjustment for multiple comparison; p values for RNAseq data are adjusted for false discovery rate. All p-values are represented as follows: * ≤ 0.05, ** ≤ 0.02, *** ≤ 0.001, **** < 0.0001.

### Bulk RNA-seq

#### Generation of bulk RNA-seq data

Naïve CD4^+^ T cells (CD4^+^CD8^neg^CD45RA^+^CD27^+^CCR7^+^CD95^neg^) were isolated by FACS from CB and AB samples. For baseline (non-activated) cells, 1 million naïve CD4^+^ T cells were washed with PBS, pelleted in an Eppendorf tube, and frozen at -80C to lyse and store cells. For the other time points, 1 million sorted naïve cells were seeded in a 96-well plate at a density of 100,000 cells/well and cultured at 37C with Immunocult^TM^ anti-CD3/CD28 soluble antibodies. Cultures were harvested at 3hr, 24hr, or 48hr post-activation and processed the same as the baseline time point. Frozen cell pellets were shipped to Novogene on dry ice, where sample quality control, library construction, and sequencing occurred according to their standard protocol^94^. Messenger RNA was purified from total RNA using poly-T oligo-attached magnetic beads. After fragmentation, the first strand cDNA was synthesized using random hexamer primers, followed by the second strand cDNA synthesis using either dUTP for directional library or DTTP for non-directional library. For the non-directional library, it was ready after end repair, A-tailing, adapter ligation, size selection, amplification, and purification. For the directional library, it was ready after end repair, A-tailing, adapter ligation, size selection, USER enzyme digestion, amplification, and purification. The library was checked with Qubit and real-time PCR for quantification and bioanalyzer for size distribution detection. Quantified libraries were pooled and sequenced on Illumina Novaseq 6000, S4 Flowcell, Paired-End 150 cycles. Samples that passed QC were advanced to RNA sequencing and included in data analysis. Six samples for each age and time point were submitted, but only those that passed RNA quality control were advanced to sequencing and analysis. The final number of individual samples from different donors for each time point are as follows: 0HR: 6CB, 5AB; 3HR: 5CB, 5AB; 24HR: 5CB, 5AB; 48HR: 5CB, 5AB.

#### Bulk RNA-seq data Processing

cDNA and ncRNA fasta files from Ensembl GRCh38 (version 109)^95^ were concatenated and used as input to the kallisto (0.46.2)^96^ ‘index’ function with default arguments. The kallisto ‘quant’ function was used to perform paired-end transcript quantification. The transcript-level estimates from kallisto were imported into R (version 4.4.1)^97^ and summarized to gene estimates using the tximport (1.32.0)^98^ package (’type = “kallisto”, countsFromAbundance = “lengthScaledTPM”, ignoreTxVersion = TRUÈ) from Bioconductor (version 3.19)^99^. Ensembl IDs were mapped to gene symbols using the ‘makeTxDbFromEnsembl’ function from the txdbmaker (1.0.1)^100^ and biomaRt packages^101^ (2.60.1) using version 109 of the ‘hsapiens_gene_ensembl’ dataset. Genes with scaffold and haplotype chromosome annotations were removed prior to downstream analysis.

These gene counts were then used to create a ‘DEGList’ object, followed by filtering on lowly expressed genes using the function ‘filterByExprs’ with the group argument set to the interaction of sample age (cord blood (CB), adult blood (AB)) and time point (0, 3, 24, 48 hours) using the edgeR (4.2.2) package^102^. Normalization factors were generated using the trimmed mean of M-values (’TMM’) method^103^. The ‘cpm’ function from edgeR with a prior count of 3 was used to generate normalized log2 counts per million (CPM) as input for principal component analysis (PCA) and heatmaps after regressing on the effect of sex using the ‘removeBatchEffect’ function from the limma (3.60.6)^104^. Gene count plots were generated with CPM values without log transformation. PCA was performed using the ‘prcomp’ function from the R stats package (4.4.1) with ‘scale = TRUÈ on the 500 most variable genes, as in the ‘plotPCÀ function from DESeq2 (1.44.0)^105^. Heatmaps were generated using 2,000 features with the highest row variances after excluding T cell receptor variable and major histocompatibility complex associated genes. Row transformation and ordering of the heatmaps was performed using the ‘coolmap’ function from limma with the argument ‘cluster.by = “de pattern”. The ‘cuttreè function from the R stats package was used to identify hierarchical clusters of genes with ‘k = 8’ clusters. Mean gene z-scores were used to summarize moving expression averages of the clusters across time points.

#### Differential expression of bulk RNA-seq

Differential gene expression and summary gene statistics were analyzed using linear models (’lmFit’) with empirical Bayes moderation (’eBayes’) and observational weighting (’voomWithQualityWeights’) with the limma package^106, 107^. A null intercept model accounting for group (interaction of age and time point) and sex was used for the ‘design’ argument. Adjusted p-values were obtained using the Benjamini-Hochberg correction using the ‘topTablè function from limma. Statistical significance was determined based on a log2 fold change threshold of 0.585 and a Benjamini-Hochberg adjusted p-value threshold of 0.05.

#### Gene set enrichment

Gene Ontology Biological Processes (GO:BP) pathways were obtained from the ontology gene set collection (C5) from the Molecular Signatures Database (MSigDB)^108, 109^. Gene Set Enrichment Analysis (GSEA) was performed using the fgsea package (1.30.0) with ‘scoreType = “std” ‘^44, 45^. Gene enrichment statistics were ranked by descending log2 fold change and genesets were required to contain at least 15 unique genes (’minSize = 15).

#### Analysis of scRNA-seq ata

Raw counts were first normalized to total counts of 1e4 and log-transformed. Then, differential expression was performed using a Wilcoxon rank-sum test provided by the sc.pp.rank_genes_groups method in the scanpy package (version 1.9.4)^110^. Genes with log-fold change > 4 or < -4 were shifted to 4 and -4 respectively for display purposes in Figure 2E. Metabolic genes were curated from the Human1 GEM^111^.

#### Compass analysis

Pseudobulking was performed on the scRNA-seq dataset by averaging gene expression profiles of single cells that belonged to the same donor (n=5 for CB and AB groups Module-Compass^30^ was performed on central carbon pathways (glycolysis, TCA, and OXPHOS) reactions curated from the Human1 GEM^111^. Raw reaction scores were negative transformed and shifted as described in [Compass reference]. Variance shrinkage and linear modeling of transformed reaction scores with condition as the independent variable were performed using the limma package^104^.

### Mass cytometry

#### Mass cytometry antibody conjugation

The mass cytometry antibodies and associated staining concentrations can be found in **Table S1**. All primary conjugations of mass cytometry antibodies were performed using Maxpar antibody labelling kits (Standard BioTools Catalog No. 201155A) according to the manufacturer’s instructions. Mass cytometry antibody conjugation quality was assessed by measuring IgG protein concentration with a NanoDrop (Thermo Fisher) and by confirming metal presence on a Helios mass cytometer (Standard BioTools). After labelling, antibody stabilization buffer (Candor Bioscience, Catalog No. 131050) supplemented with 0.1% sodium azide was added to generate final antibody concentrations in the range 0.2 – 0.5 mg/mL. All antibodies were stored long term at 4°C.

#### Mass cytometry barcoding and staining

Cells from five adult donors and five cord blood donors were thawed and washed before resting overnight in a 37°C 5% CO2 incubator in Resting Media (RPMI with 10% FBS, 1% 100x Pen/Strep, 1% Hepes, and 1% Glutamax). For cell viability measurement, 2.5 million cells per donor were re-suspended in PBS + 5mM EDTA (Standard BioTools, Catalog No. 201058) and incubated for exactly 60 seconds with 25 μM Cisplatin (Sigma-Aldrich, Catalog No. P4394). Cells were washed once with cell staining buffer (Standard BioTools, Catalog No. 201068) and fixed with 1.6% paraformaldehyde (Electron Microscopy Sciences, Catalog No. 15710) for 10 minutes at room temperature. For all wash steps, cells were spun at 600 x g for 5 minutes at 4°C. Cells were then barcoded using the Cell-ID 20-Plex Palladium Barcoding Kit (Standard BioTools, Catalog No. 201060) by incubating a unique barcode with each sample in Maxpar Barcode Perm Buffer for 20 minutes at room temperature on a shaker. Surface and intracellular antibody staining cocktails were prepared ahead of time in cell staining buffer or Foxp3 staining kit buffer (eBioscience, Catalog No. 00-5523-00) respectively and filtered through 0.1-μm spin-column (Millipore, Catalog No. UFC30VV00) to remove antibody aggregates. The surface antibody cocktail included Human TruStain FcX ™ at 20 mg/mL (BioLegend, Catalog No. 422302). Combined barcoded samples were incubated with 500 μL of surface antibody staining cocktail for 30 minutes at room temperature on a shaker. Cells were then washed two times with cell staining buffer before a second 1.6% PFA fix for ten minutes at room temperature. Following a single wash with cell staining buffer, cells were permeabilized with 300 μL of ice cold 100% methanol for 10 minutes at 4°C with gentle pipetting to mix halfway through. Cells were washed once with cell staining buffer and twice with Foxp3 staining kit buffer to ensure all methanol was removed. A total volume of 500 μL of intracellular antibody cocktail was incubated with the cells for 60 minutes at room temperature on a shaker. The intracellular antibody cocktail was removed with two washes of Foxp3 staining kit buffer and one wash of cell staining buffer. Cells were re-suspended overnight in PBS (Standard BioTools, Catalog No. 201058) containing 4% PFA and 1:1,500 191/193Ir Cell-ID Intercalator Solution (Standard BioTools, Catalog No. 201192A).

#### Mass cytometry data acquisition

Prior to running mass cytometry, cells were washed once with cell staining media, once with PBS and once with MilliQ water. Cells were filtered through a filter cap tube (Fisher Scientific, Catalog No. 352235) and resuspended at a concentration of 1x10^6 cells/mL in MilliQ water containing a 1:50 ratio of EQ four element calibration beads (Standard BioTools, Catalog No. 201078). Samples were acquired at an event rate of 400 – 600 events/second on a Helios mass cytometer (Standard BioTools).

#### Mass cytometry data analysis

Bead based data normalization and de-barcoding of combined samples was performed using the R package Premessa available at https://github.com/ParkerICI/premessa. All analyses were performed on naïve CD4^+^ T cells (CD45RA^+^CCR7^+^) manually gated in CellEngine (CellCarta Fremont LLC) and exported as FCS files before uploading to R using flowCore^112^. Marker expression was arcsinh transformed using a cofactor of 5 for visualizations. The CATALYST R/Bioconductor package^54^ and FlowSOM^53^ clustering algorithm were used to generate clusters based on T cell and metabolic state markers (CD98, CD69, LDH, PFKFB4, CS, ACADM, GLUD1, ATP5a, VDAC1, FoxP3, GLUT1, CytC, GAPDH, ASCT2, CPT1a, MCT1, CD38, FOXP3). Initial clustering yielded 10 clusters that were manually merged to 8 clusters based on marker expression. UMAPs were used to visualize clusters and marker expression across clusters in combination with heatmaps showing marker expression levels. For all clusters, we used the R package diffcyt^113^ to perform differential abundance analysis using a generalized linear model with age (CB, AB) as a fixed effect and donor as a random effect.

### Data Availability and Software Used

- Schematics were made with Biorender (https://biorender.com). GraphPad (Prism) 10 was used. R version 4.4.1 was used for bulk and single cell RNA-seq analyses.
- The bulk RNA-seq data generated in this study has been deposited to the GEO as indicated in the key resources table.
- Any additional information required to reanalyze the data reported in this paper is available from the lead contact upon request.

## References

1. Gensollen, T., and Blumberg, R.S. (2017). Correlation between early-life regulation of the immune system by microbiota and allergy development. Journal of Allergy and Clinical Immunology 139, 1084–1091. 10.1016/J.JACI.2017.02.011.

2. Yang, S., Fujikado, N., Kolodin, D., Benoist, C., and Mathis, D. (2015). Immune tolerance. Regulatory T cells generated early in life play a distinct role in maintaining self-tolerance. Science 348, 589–594. 10.1126/SCIENCE.AAA7017.

3. Suri-Payer, E., Amar, A.Z., Thornton, A.M., and Shevach, E.M. (1998). CD4+CD25+ T cells inhibit both the induction and effector function of autoreactive T cells and represent a unique lineage of immunoregulatory cells. J Immunol 160, 1212–1218. 10.4049/jimmunol.160.3.1212.

4. Asano, M., Toda, M., Sakaguchi, N., and Sakaguchi, S. (1996). Autoimmune disease as a consequence of developmental abnormality of a T cell subpopulation. J Exp Med 184, 387 – 396.

5. Wang, G., Miyahara, Y., Guo, Z., Khattar, M., Stepkowski, S.M., and Chen, W. (2010). “Default” generation of neonatal regulatory T cells. J Immunol 185, 71–78. 10.4049/JIMMUNOL.0903806.

6. Mold, J.E., Venkatasubrahmanyam, S., Burt, T.D., Michaëlsson, J., Rivera, J.M., Galkina, S.A., Weinberg, K., Stoddart, C.A., and McCune, J.M. (2010). Fetal and Adult Hematopoietic Stem Cells Give Rise to Distinct T Cell Lineages in Humans. Science (1979) 330, 1695–1699. 10.1126/SCIENCE.1196509.

7. Ng, M.S.F., Roth, T.L., Mendoza, V.F., Marson, A., and Burt, T.D. (2019). Helios enhances the preferential differentiation of human fetal CD4+ naïve T cells into regulatory T cells. Sci Immunol 4. 10.1126/SCIIMMUNOL.AAV5947.

8. Bunis, D.G., Bronevetsky, Y., Krow-Lucal, E., Bhakta, N.R., Kim, C.C., Nerella, S., Jones, N., Mendoza, V.F., Bryson, Y.J., Gern, J.E., et al. (2021). Single-Cell Mapping of Progressive Fetal-to-Adult Transition in Human Naive T Cells. Cell Rep 34. 10.1016/J.CELREP.2020.108573.

9. Scharschmidt, T.C., Vasquez, K.S., Truong, H.-A., Gearty, S. V, Pauli, M.L., Nosbaum, A., Gratz, I.K., Otto, M., Moon, J.J., Liese, J., et al. (2015). A wave of regulatory T cells into neonatal skin mediates tolerance to commensal microbes. Immunity 43.

10. Akagbosu, B., Tayyebi, Z., Shibu, G., Paucar Iza, Y.A., Deep, D., Parisotto, Y.F., Fisher, L., Pasolli, H.A., Thevin, V., Elmentaite, R., et al. (2022). Novel antigen-presenting cell imparts Treg-dependent tolerance to gut microbiota. Nature 610, 752–760. 10.1038/S41586-022-05309-5.

11. Thome, J.J.C., Bickham, K.L., Ohmura, Y., Kubota, M., Matsuoka, N., Gordon, C., Granot, T., Griesemer, A., Lerner, H., Kato, T., et al. (2016). Early-life compartmentalization of human T cell differentiation and regulatory function in mucosal and lymphoid tissues. Nat Med 22, 72–77. 10.1038/NM.4008.

12. Toit, G. Du, Roberts, G., Sayre, P.H., Bahnson, H.T., Radulovic, S., Santos, A.F., Brough, H.A., Phippard, D., Basting, M., Feeney, M., et al. (2015). Randomized trial of peanut consumption in infants at risk for peanut allergy. N Engl J Med 372, 803 – 813. 10.1056/nejmoa1414850.

13. Thornton, C.A., Upham, J.W., Wikström, M.E., Holt, B.J., White, G.P., Sharp, M.J., Sly, P.D., and Holt, P.G. (2004). Functional Maturation of CD4+CD25+CTLA4+CD45RA+ T Regulatory Cells in Human Neonatal T Cell Responses to Environmental Antigens/Allergens. The Journal of Immunology 173, 3084–3092. 10.4049/JIMMUNOL.173.5.3084.

14. Anderson, K.A., Madsen, A.S., Olsen, C.A., and Hirschey, M.D. (2017). Metabolic control by sirtuins and other enzymes that sense NAD+, NADH, or their ratio. Biochimica et Biophysica Acta (BBA) - Bioenergetics 1858, 991–998. 10.1016/J.BBABIO.2017.09.005.

15. Wu, H., Huang, H., and Zhao, Y. (2023). Interplay between metabolic reprogramming and post-translational modifications: from glycolysis to lactylation. Front Immunol 14. 10.3389/FIMMU.2023.1211221.

16. Holm, S.R., Jenkins, B.J., Cronin, J.G., Jones, N., and Thornton, C.A. (2021). A role for metabolism in determining neonatal immune function. Pediatric Allergy and Immunology 32, 1616–1628. 10.1111/PAI.13583.

17. Pearce, E.L. (2010). Metabolism in T cell activation and differentiation. Curr Opin Immunol 22, 314. 10.1016/J.COI.2010.01.018.

18. Chapman, N.M., Boothby, M.R., and Chi, H. (2019). Metabolic coordination of T cell quiescence and activation. Nature Reviews Immunology 2019 20:1 20, 55–70. 10.1038/s41577-019-0203-y.

19. Tabilas, C., Wang, J., Liu, X., Locasale, J.W., Smith, N.L., and Rudd, B.D. (2019). Cutting Edge: Elevated Glycolytic Metabolism Limits the Formation of Memory CD8+ T Cells in Early Life. The Journal of Immunology 203, 2571–2576. 10.4049/JIMMUNOL.1900426.

20. Ghosh, A., Khanam, A., Ray, K., Mathur, P., Subramanian, A., Poonia, B., and Kottilil, S. (2023). CD38: an ecto-enzyme with functional diversity in T cells. Front Immunol 14, 1146791. 10.3389/FIMMU.2023.1146791.

21. Kar, A., Mehrotra, S., and Chatterjee, S. (2020). CD38: T Cell Immuno-Metabolic Modulator. Cells 2020, Vol. 9, Page 1716 9, 1716. 10.3390/CELLS9071716.

22. DeRogatis, J.M., Neubert, E.N., Viramontes, K.M., Henriquez, M.L., Nicholas, D.A., and Tinoco, R. (2023). Cell-Intrinsic CD38 Expression Sustains Exhausted CD8+ T Cells by Regulating Their Survival and Metabolism during Chronic Viral Infection. J Virol 97. 10.1128/JVI.00225-23.

23. Chatterjee, S., Daenthanasanmak, A., Chakraborty, P., Wyatt, M.W., Dhar, P., Selvam, S.P., Fu, J., Zhang, J., Nguyen, H., Kang, I., et al. (2018). CD38-NAD + Axis Regulates Immunotherapeutic Anti-Tumor T Cell Response. Cell Metab 27, 85–100.e8. 10.1016/J.CMET.2017.10.006/ATTACHMENT/085FD262-CFBA-4054-B132-E23D1F4E88CC/MMC2.PDF.

24. Reolo, M.J.Y., Otsuka, M., Seow, J.J.W., Lee, J., Lee, Y.H., Nguyen, P.H.D., Lim, C.J., Wasser, M., Chua, C., Lim, T.K.H., et al. (2023). CD38 marks the exhausted CD8+ tissue-resident memory T cells in hepatocellular carcinoma. Front Immunol 14. 10.3389/FIMMU.2023.1182016.

25. Bohacova, P., Terekhova, M., Tsurinov, P., Mullins, R., Husarcikova, K., Shchukina, I., Antonova, A.U., Echalar, B., Kossl, J., Saidu, A., et al. (2024). Multidimensional profiling of human T cells reveals high CD38 expression, marking recent thymic emigrants and age-related naive T cell remodeling. Immunity 57, 2362–2379. 10.1016/J.IMMUNI.2024.08.019.

26. Piedra-Quintero, Z.L., Wilson, Z., Nava, P., and Guerau-de-Arellano, M. (2020). CD38: An Immunomodulatory Molecule in Inflammation and Autoimmunity. Front Immunol 11, 597959. 10.3389/FIMMU.2020.597959.

27. Wei, W., Graeff, R., and Yue, J. (2014). Roles and mechanisms of the CD38/cyclic adenosine diphosphate ribose/Ca2+ signaling pathway. World J Biol Chem 5, 58. 10.4331/WJBC.V5.I1.58.

28. Cho, Y.S., Han, M.K., Choi, Y.B., Yun, Y., Shin, J., and Kim, U.H. (2000). Direct interaction of the CD38 cytoplasmic tail and the Lck SH2 domain. Cd38 transduces T cell activation signals through associated Lck. J Biol Chem 275, 1685–1690. 10.1074/JBC.275.3.1685.

29. Ghosh, A., Khanam, A., Ray, K., Mathur, P., Subramanian, A., Poonia, B., and Kottilil, S. (2023). CD38: an ecto-enzyme with functional diversity in T cells. Front Immunol 14, 1146791. 10.3389/FIMMU.2023.1146791.

30. Wagner, A., Wang, C., Fessler, J., DeTomaso, D., Avila-Pacheco, J., Kaminski, J., Zaghouani, S., Christian, E., Thakore, P., Schellhaass, B., et al. (2021). Metabolic modeling of single Th17 cells reveals regulators of autoimmunity. Cell 184, 4168–4185.e21. 10.1016/J.CELL.2021.05.045.

31. Li, W., Liang, L., Liao, Q., Li, Y., and Zhou, Y. (2022). CD38: An important regulator of T cell function. Biomedicine and Pharmacotherapy 153. 10.1016/J.BIOPHA.2022.113395.

32. Van Loosdregt, J., Vercoulen, Y., Guichelaar, T., Gent, Y.Y.J., Beekman, J.M., Van Beekum, O., Brenkman, A.B., Hijnen, D.J., Mutis, T., Kalkhoven, E., et al. (2010). Regulation of Treg functionality by acetylation-mediated Foxp3 protein stabilization. Blood 115, 965–974. 10.1182/BLOOD-2009-02-207118.

33. Loosdregt, J. van, Brunen, D., Fleskens, V., Pals, C.E.G.M., Lam, E.W.F., and Coffer, P.J. (2011). Rapid Temporal Control of Foxp3 Protein Degradation by Sirtuin-1. PLoS One 6, e19047. 10.1371/JOURNAL.PONE.0019047.

34. Sequeira, J., Boily, G., Bazinet, S., Saliba, S., He, X., Jardine, K., Kennedy, C., Staines, W., Rousseaux, C., Mueller, R., et al. (2008). sirt1-null mice develop an autoimmune-like condition. Exp Cell Res 314, 3069–3074. 10.1016/J.YEXCR.2008.07.011.

35. Zhang, J., Lee, S.M., Shannon, S., Gao, B., Chen, W., Chen, A., Divekar, R., McBurney, M.W., Braley-Mullen, H., Zaghouani, H., et al. (2009). The type III histone deacetylase Sirt1 is essential for maintenance of T cell tolerance in mice. J Clin Invest 119, 3048. 10.1172/JCI38902.

36. Tabilas, C., Wang, J., Liu, X., Locasale, J.W., Smith, N.L., and Rudd, B.D. (2019). Cutting Edge: Elevated Glycolytic Metabolism Limits the Formation of Memory CD8+ T Cells in Early Life. The Journal of Immunology 203, 2571–2576. 10.4049/JIMMUNOL.1900426.

37. Zhang, Y., Maksimovic, J., Huang, B., De Souza, D.P., Naselli, G., Chen, H., Zhang, L., Weng, K., Liang, H., Xu, Y., et al. (2018). Cord Blood CD8+ T Cells Have a Natural Propensity to Express IL-4 in a Fatty Acid Metabolism and Caspase Activation-Dependent Manner. Front Immunol 9, 358166. 10.3389/FIMMU.2018.00879.

38. Kar, A., Mehrotra, S., and Chatterjee, S. (2020). CD38: T Cell Immuno-Metabolic Modulator. Cells 2020, Vol. 9, Page 1716 *9*, 1716. 10.3390/CELLS9071716.

39. Rudd, B.D. (2020). Neonatal T Cells: A Reinterpretation. Annu Rev Immunol 38, 229–247. 10.1146/ANNUREV-IMMUNOL-091319-083608.

40. Bohacova, P., Terekhova, M., Tsurinov, P., Mullins, R., Husarcikova, K., Shchukina, I., Antonova, A.U., Echalar, B., Kossl, J., Saidu, A., et al. (2024). Multidimensional profiling of human T cells reveals high CD38 expression, marking recent thymic emigrants and age-related naive T cell remodeling. Immunity 57, 2362–2379.e10. 10.1016/J.IMMUNI.2024.08.019.

41. Kimmig, S., Przybylski, G.K., Schmidt, C.A., Laurisch, K., Möwes, B., Radbruch, A., and Thiel, A. (2002). Two Subsets of Naive T Helper Cells with Distinct T Cell Receptor Excision Circle Content in Human Adult Peripheral Blood. Journal of Experimental Medicine 195, 789–794. 10.1084/JEM.20011756.

42. Schilham, M.W., Moerer, P., Cumano, A., and Clevers, H.C. (1997). Sox-4 facilitates thymocyte differentiation. Eur J Immunol 27, 1292–1295. 10.1002/EJI.1830270534.

43. Yu, Y., Wang, J., Khaled, W., Burke, S., Li, P., Chen, X., Yang, W., Jenkins, N.A., Copeland, N.G., Zhang, S., et al. (2012). Bcl11a is essential for lymphoid development and negatively regulates p53. J Exp Med 209, 2467. 10.1084/JEM.20121846.

44. Korotkevich, G., Sukhov, V., Budin, N., Shpak, B., Artyomov, M.N., and Sergushichev, A. (2021). Fast gene set enrichment analysis. bioRxiv, 060012. 10.1101/060012.

45. Subramanian, A., Tamayo, P., Mootha, V.K., Mukherjee, S., Ebert, B.L., Gillette, M.A., Paulovich, A., Pomeroy, S.L., Golub, T.R., Lander, E.S., et al. (2005). Gene set enrichment analysis: a knowledge-based approach for interpreting genome-wide expression profiles. Proc Natl Acad Sci U S A 102, 15545–15550. 10.1073/PNAS.0506580102.

46. Tabilas, C., Smith, N.L., and Rudd, B.D. (2023). Shaping immunity for life: Layered development of CD8+ T cells. Immunol Rev 315, 108–125. 10.1111/IMR.13185.

47. Li, X., Liang, Y., Leblanc, M., Benner, C., and Zheng, Y. (2014). Function of a Foxp3 cis-element in protecting regulatory T cell identity. Cell 158, 734. 10.1016/J.CELL.2014.07.030.

48. Feng, Y., Arvey, A., Chinen, T., Veeken, J. van der, Gasteiger, G., and Rudensky, A.Y. (2014). Control of the inheritance of regulatory T cell identity by a cis element in the Foxp3 locus. Cell 158, 749. 10.1016/J.CELL.2014.07.031.

49. Chen, K.Y., Kibayashi, T., Giguelay, A., Hata, M., Nakajima, S., Mikami, N., Takeshima, Y., Ichiyama, K., Omiya, R., Ludwig, L.S., et al. (2025). Genome-wide CRISPR screen in human T cells reveals regulators of FOXP3. Nature 2025, 1–10. 10.1038/s41586-025-08795-5.

50. Phan, A.T., Doedens, A.L., Palazon, A., Tyrakis, P.A., Cheung, K.P., Johnson, R.S., and Goldrath, A.W. (2016). Constitutive Glycolytic Metabolism Supports CD8+ T Cell Effector Memory Differentiation during Viral Infection. Immunity 45, 1024–1037. 10.1016/J.IMMUNI.2016.10.017.

51. Hartmann, F.J., Mrdjen, D., McCaffrey, E., Glass, D.R., Greenwald, N.F., Bharadwaj, A., Khair, Z., Verberk, S.G.S., Baranski, A., Baskar, R., et al. (2020). Single-cell metabolic profiling of human cytotoxic T cells. Nature Biotechnology 2020 39:2 39, 186–197. 10.1038/s41587-020-0651-8.

52. Levine, L.S., Hiam-Galvez, K.J., Marquez, D.M., Tenvooren, I., Madden, M.Z., Contreras, D.C., Dahunsi, D.O., Irish, J.M., Oluwole, O.O., Rathmell, J.C., et al. (2021). Single-cell analysis by mass cytometry reveals metabolic states of early-activated CD8+ T cells during the primary immune response. Immunity 54, 829–844.e5. 10.1016/J.IMMUNI.2021.02.018.

53. Quintelier, K., Couckuyt, A., Emmaneel, A., Aerts, J., Saeys, Y., and Van Gassen, S. (2021). Analyzing high-dimensional cytometry data using FlowSOM. Nature Protocols 2021 16:8 16, 3775–3801. 10.1038/s41596-021-00550-0.

54. Nowicka, M., Krieg, C., Crowell, H.L., Weber, L.M., Hartmann, F.J., Guglietta, S., Becher, B., Levesque, M.P., and Robinson, M.D. (2017). CyTOF workflow: differential discovery in high-throughput high-dimensional cytometry datasets. F1000Res 6, 748. 10.12688/f1000research.11622.3.

55. Zhang, Y.T., Xing, M.L., Fang, H.H., Li, W.D., Wu, L., and Chen, Z.P. (2023). Effects of lactate on metabolism and differentiation of CD4+T cells. Mol Immunol 154, 96–107. 10.1016/J.MOLIMM.2022.12.015.

56. Shyer, J.A., Flavell, R.A., and Bailis, W. (2020). Metabolic signaling in T cells. Cell Research 2020 30:8 *30*, 649–659. 10.1038/s41422-020-0379-5.

57. Qu, Q., Zeng, F., Liu, X., Wang, Q.J., and Deng, F. (2016). Fatty acid oxidation and carnitine palmitoyltransferase I: emerging therapeutic targets in cancer. Cell Death & Disease 2016 7:5 7, e2226–e2226. 10.1038/cddis.2016.132.

58. Ren, W., Liu, G., Yin, J., Tan, B., Wu, G., Bazer, F.W., Peng, Y., and Yin, Y. (2017). Amino-acid transporters in T-cell activation and differentiation. Cell Death & Disease 2017 8:3 8, e2655–e2655. 10.1038/cddis.2016.222.

59. Jin, J., Byun, J.K., Choi, Y.K., and Park, K.G. (2023). Targeting glutamine metabolism as a therapeutic strategy for cancer. Experimental & Molecular Medicine 2023 55:4 55, 706–715. 10.1038/s12276-023-00971-9.

60. Argüello, R.J., Combes, A.J., Char, R., Gigan, J.-P., Baaziz, A.I., Bousiquot, E., Camosseto, V., Samad, B., Tsui, J., Yan, P., et al. (2020). SCENITH: A Flow Cytometry-Based Method to Functionally Profile Energy Metabolism with Single-Cell Resolution. Cell Metab 32, 1063–1075.e7. 10.1016/j.cmet.2020.11.007.

61. Yagi, Y., Kuwahara, M., Suzuki, J., Imai, Y., and Yamashita, M. (2020). Glycolysis and subsequent mevalonate biosynthesis play an important role in Th2 cell differentiation. Biochem Biophys Res Commun 530, 355–361. 10.1016/J.BBRC.2020.08.009.

62. Yang, J.Q., Kalim, K.W., Li, Y., Zhang, S., Hinge, A., Filippi, M.D., Zheng, Y., and Guo, F. (2015). RhoA orchestrates glycolysis for Th2 cell differentiation and allergic airway inflammation. J Allergy Clin Immunol 137, 231. 10.1016/J.JACI.2015.05.004.

63. Fang, J., Chen, W., Hou, P., Liu, Z., Zuo, M., Liu, S., Feng, C., Han, Y., Li, P., Shi, Y., et al. (2023). NAD+ metabolism-based immunoregulation and therapeutic potential. Cell & Bioscience 2023 13:1 *13*, 1–15. 10.1186/S13578-023-01031-5.

64. Jeng, M.Y., Hull, P.A., Fei, M., Kwon, H.-S., Tsou, C.-L., Kasler, H., Ng, C.-P., Gordon, D.E., Johnson, J., Krogan, N., et al. (2018). Metabolic reprogramming of human CD8+ memory T cells through loss of SIRT1. J Exp Med 215, 51. 10.1084/JEM.20161066.

65. Zhang, X., Xiao, X., Lan, P., Li, J., Dou, Y., Chen, W., Ishii, N., Chen, S., Xia, B., Chen, K., et al. (2018). OX40 Costimulation Inhibits Foxp3 Expression and Treg Induction via BATF3-Dependent and Independent Mechanisms. Cell Rep 24, 607–618. 10.1016/J.CELREP.2018.06.052.

66. Li, J., Xu, B., He, M., Zong, X., Cunningham, T., Sha, C., Fan, Y., Cross, R., Hanna, J.H., and Feng, Y. (2021). Control of Foxp3 induction and maintenance by sequential histone acetylation and DNA demethylation. Cell Rep 37, 110124. 10.1016/J.CELREP.2021.110124.

67. Mayoral, R., Osborn, O., McNelis, J., Johnson, A.M., Oh, D.Y., Izquierdo, C.L., Chung, H., Li, P., Traves, P.G., Bandyopadhyay, G., et al. (2015). Adipocyte SIRT1 knockout promotes PPARγ activity, adipogenesis and insulin sensitivity in chronic-HFD and obesity. Mol Metab 4, 378–391. 10.1016/J.MOLMET.2015.02.007.

68. Han, L., Zhou, R., Niu, J., McNutt, M.A., Wang, P., and Tong, T. (2010). SIRT1 is regulated by a PPARγ–SIRT1 negative feedback loop associated with senescence. Nucleic Acids Res 38, 7458. 10.1093/NAR/GKQ609.

69. Lehrke, M., and Lazar, M.A. (2005). The many faces of PPARγ. Cell 123, 993–999. 10.1016/J.CELL.2005.11.026/ASSET/75EFF205-2B62-4C1D-A347-BA858A6E027B/MAIN.ASSETS/GR1.JPG.

70. Cipolletta, D., Feuerer, M., Li, A., Kamei, N., Lee, J., Shoelson, S.E., Benoist, C., and Mathis, D. (2012). PPARγ is a major driver of the accumulation and phenotype of adipose-tissue Treg cells. Nature 486, 549. 10.1038/NATURE11132.

71. Hontecillas, R., and Bassaganya-Riera, J. (2007). Peroxisome Proliferator-Activated Receptor γ Is Required for Regulatory CD4+ T Cell-Mediated Protection against Colitis. The Journal of Immunology 178, 2940–2949. 10.4049/JIMMUNOL.178.5.2940.

72. Guri, A.J., Mohapatra, S.K., Horne, W.T., Hontecillas, R., and Bassaganya-Riera, J. (2010). The role of T cell PPAR gamma in mice with experimental inflammatory bowel disease. BMC Gastroenterol 10. 10.1186/1471-230X-10-60.

73. Wohlfert, E.A., Nichols, F.C., Nevius, E., and Clark, R.B. (2007). Peroxisome Proliferator-Activated Receptor γ (PPARγ) and Immunoregulation: Enhancement of Regulatory T Cells through PPARγ-Dependent and -Independent Mechanisms. The Journal of Immunology 178, 4129–4135. 10.4049/JIMMUNOL.178.7.4129.

74. Ferreira, L.M.R., Muller, Y.D., Bluestone, J.A., and Tang, Q. (2019). Next-generation regulatory T cell therapy. Nat Rev Drug Discov 18, 749–769. 10.1038/S41573-019-0041-4.

75. Wang, Q., Hu, J., Han, G., Wang, P., Li, S., Chang, J., Gao, K., Yin, R., Li, Y., Zhang, T., et al. (2022). PTIP governs NAD+ metabolism by regulating CD38 expression to drive macrophage inflammation. Cell Rep 38. 10.1016/J.CELREP.2022.110603.

76. Callen, E., Faryabi, R.B., Luckey, M., Hao, B., Daniel, J.A., Yang, W., Sun, H.W., Dressler, G., Peng, W., Chi, H., et al. (2012). The DNA damage- and transcription-associated protein paxip1 controls thymocyte development and emigration. Immunity 37, 971–985. 10.1016/J.IMMUNI.2012.10.007.

77. Dong, M., Mallet Gauthier, È., Fournier, M., and Melichar, H.J. (2022). Developing the right tools for the job: Lin28 regulation of early life T-cell development and function. FEBS J 289, 4416–4429. 10.1111/FEBS.16045.

78. Zhu, H., Ng, S.C., Segr, A. V., Shinoda, G., Shah, S.P., Einhorn, W.S., Takeuchi, A., Engreitz, J.M., Hagan, J.P., Kharas, M.G., et al. (2011). The Lin28/let-7 axis regulates glucose metabolism. Cell 147, 81–94. 10.1016/J.CELL.2011.08.033.

79. Murray, J.M., Kaufmann, G.R., Hodgkin, P.D., Lewin, S.R., Kelleher, A.D., Davenport, M.P., and Zaunders, J.J. (2003). Naive T cells are maintained by thymic output in early ages but by proliferation without phenotypic change after age twenty. Immunol Cell Biol 81, 487–495. 10.1046/J.1440-1711.2003.01191.X.

80. Holm, S.R., Jenkins, B.J., Cronin, J.G., Jones, N., and Thornton, C.A. (2021). A role for metabolism in determining neonatal immune function. Pediatric Allergy and Immunology 32, 1616–1628. 10.1111/PAI.13583.

81. Sánchez-Villanueva, J.A., Rodríguez-Jorge, O., Ramírez-Pliego, O., Salgado, G.R., Abou-Jaoudé, W., Hernandez, C., Naldi, A., Thieffry, D., and Santana, M.A. (2019). Contribution of ROS and metabolic status to neonatal and adult CD8+ T cell activation. PLoS One 14, e0226388. 10.1371/JOURNAL.PONE.0226388.

82. De Rosa, V., Galgani, M., Porcellini, A., Colamatteo, A., Santopaolo, M., Zuchegna, C., Romano, A., De Simone, S., Procaccini, C., La Rocca, C., et al. (2015). Glycolysis controls the induction of human regulatory T cells by modulating the expression of FOXP3 exon 2 splicing variants. Nature Immunology 2015 16:11 16, 1174–1184. 10.1038/ni.3269.

83. Du, J., Wang, Q., Yang, S., Chen, S., Fu, Y., Spath, S., Domeier, P., Hagin, D., Anover-Sombke, S., Haouili, M., et al. (2022). FOXP3 exon 2 controls Treg stability and autoimmunity. Sci Immunol 7. 10.1126/SCIIMMUNOL.ABO5407.

84. Kempkes, R.W.M., Joosten, I., Koenen, H.J.P.M., and He, X. (2019). Metabolic Pathways Involved in Regulatory T Cell Functionality. Front Immunol 10, 483290. 10.3389/FIMMU.2019.02839.

85. Tomaszewicz, M., Ronowska, A., Zieliński, M., Jankowska-Kulawy, A., and Trzonkowski, P. (2023). T regulatory cells metabolism: The influence on functional properties and treatment potential. Front Immunol 14, 1122063. 10.3389/FIMMU.2023.1122063/BIBTEX.

86. Rogina, B., and Tissenbaum, H.A. (2024). SIRT1, resveratrol and aging. Front Genet 15, 1393181. 10.3389/FGENE.2024.1393181/BIBTEX.

87. Yang Xiao, Lun Yu, Jiang Han, Liu Xun, Duan Zhiquan, Xin Shijie, and Zhang Jian (2018). SIRT1-Regulated Abnormal Acetylation of FOXP3 Induces Regulatory T-Cell Function Defect in Hashimoto’s Thyroiditis. https://home.liebertpub.com/thy 28, 246–256. 10.1089/THY.2017.0286.

88. Rodgers, J.T., Lerin, C., Haas, W., Gygi, S.P., Spiegelman, B.M., and Puigserver, P. (2005). Nutrient control of glucose homeostasis through a complex of PGC-1α and SIRT1. Nature 2005 434:7029 434, 113–118. 10.1038/nature03354.

89. Lim, J.H., Lee, Y.M., Chun, Y.S., Chen, J., Kim, J.E., and Park, J.W. (2010). Sirtuin 1 Modulates Cellular Responses to Hypoxia by Deacetylating Hypoxia-Inducible Factor 1α. Mol Cell 38, 864–878. 10.1016/J.MOLCEL.2010.05.023.

90. Hallows, W.C., Yu, W., and Denu, J.M. (2012). Regulation of Glycolytic Enzyme Phosphoglycerate Mutase-1 by Sirt1 Protein-mediated Deacetylation. Journal of Biological Chemistry 287, 3850–3858. 10.1074/JBC.M111.317404.

91. Bertschi, N.L., Steck, O., Luther, F., Bazzini, C., von Meyenn, L., Schärli, S., Vallone, A., Felser, A., Keller, I., Friedli, O., et al. (2023). PPAR-γ regulates the effector function of human T helper 9 cells by promoting glycolysis. Nature Communications 2023 14:1 14, 1–13. 10.1038/s41467-023-38233-x.

92. Ng, M.S.F., Roth, T.L., Mendoza, V.F., Marson, A., and Burt, T.D. (2019). Helios enhances the preferential differentiation of human fetal CD4+ naïve T cells into regulatory T cells. Sci Immunol 4. 10.1126/SCIIMMUNOL.AAV5947.

93. Kressler, C., Gasparoni, G., Nordström, K., Hamo, D., Salhab, A., Dimitropoulos, C., Tierling, S., Reinke, P., Volk, H.D., Walter, J., et al. (2021). Targeted De-Methylation of the FOXP3-TSDR Is Sufficient to Induce Physiological FOXP3 Expression but Not a Functional Treg Phenotype. Front Immunol 11. 10.3389/FIMMU.2020.609891.

94. Parkhomchuk, D., Borodina, T., Amstislavskiy, V., Banaru, M., Hallen, L., Krobitsch, S., Lehrach, H., and Soldatov, A. (2009). Transcriptome analysis by strand-specific sequencing of complementary DNA. Nucleic Acids Res 37, e123. 10.1093/NAR/GKP596.

95. Martin, F.J., Amode, M.R., Aneja, A., Austine-Orimoloye, O., Azov, A.G., Barnes, I., Becker, A., Bennett, R., Berry, A., Bhai, J., et al. (2023). Ensembl 2023. Nucleic Acids Res 51, D933–D941. 10.1093/NAR/GKAC958.

96. Bray, N.L., Pimentel, H., Melsted, P., and Pachter, L. (2016). Near-optimal probabilistic RNA-seq quantification. Nature Biotechnology 2016 34:5 34, 525–527. 10.1038/nbt.3519.

97. R Core Team (2024) (2024). R: A Language and Environment for Statistical Computing. Preprint at R Foundation for Statistical Computing.

98. Soneson, C., Love, M.I., and Robinson, M.D. (2016). Differential analyses for RNA-seq: transcript-level estimates improve gene-level inferences. F1000Research 2016 4:1521 4, 1521. 10.12688/f1000research.7563.2.

99. Huber, W., Carey, V.J., Gentleman, R., Anders, S., Carlson, M., Carvalho, B.S., Bravo, H.C., Davis, S., Gatto, L., Girke, T., et al. (2015). Orchestrating high-throughput genomic analysis with Bioconductor. Nature Methods 2015 12:2 12, 115–121. 10.1038/nmeth.3252.

100. Pagès H, C.M.A.P.F.S.M.M. (2024). txdbmaker: Tools for making TxDb objects from genomic annotations. Preprint.

101. Durinck, S., Spellman, P.T., Birney, E., and Huber, W. (2009). Mapping identifiers for the integration of genomic datasets with the R/Bioconductor package biomaRt. Nat Protoc 4, 1184–1191. 10.1038/NPROT.2009.97.

102. Chen, Y., Chen, L., Lun, A.T.L., Baldoni, P.L., and Smyth, G.K. (2024). edgeR v4: powerful differential analysis of sequencing data with expanded functionality and improved support for small counts and larger datasets. bioRxiv, 2024.01.21.576131. 10.1101/2024.01.21.576131.

103. Robinson, M.D., and Oshlack, A. (2010). A scaling normalization method for differential expression analysis of RNA-seq data. Genome Biol 11, 1–9. 10.1186/GB-2010-11-3-R25/FIGURES/3.

104. Ritchie, M.E., Phipson, B., Wu, D., Hu, Y., Law, C.W., Shi, W., and Smyth, G.K. (2015). limma powers differential expression analyses for RNA-sequencing and microarray studies. Nucleic Acids Res 43, e47–e47. 10.1093/NAR/GKV007.

105. Love, M.I., Huber, W., and Anders, S. (2014). Moderated estimation of fold change and dispersion for RNA-seq data with DESeq2. Genome Biology 2014 15:12 15, 1–21. 10.1186/S13059-014-0550-8.

106. Phipson, B., Lee, S., Majewski, I.J., Alexander, W.S., and Smyth, G.K. (2016). Robust hyperparameter estimation protects against hypervariable genes and improves power to detect differential expression. 10.1214/16-AOAS920 10, 946–963. 10.1214/16-AOAS920.

107. Liu, R., Holik, A.Z., Su, S., Jansz, N., Chen, K., Leong, H.S., Blewitt, M.E., Asselin-Labat, M.L., Smyth, G.K., and Ritchie, M.E. (2015). Why weight? Modelling sample and observational level variability improves power in RNA-seq analyses. Nucleic Acids Res 43, e97. 10.1093/NAR/GKV412.

108. Consortium, T.G.O., Aleksander, S.A., Balhoff, J., Carbon, S., Cherry, J.M., Drabkin, H.J., Ebert, D., Feuermann, M., Gaudet, P., Harris, N.L., et al. (2023). The Gene Ontology knowledgebase in 2023. Genetics 224. 10.1093/GENETICS/IYAD031.

109. Ashburner, M., Ball, C.A., Blake, J.A., Botstein, D., Butler, H., Cherry, J.M., Davis, A.P., Dolinski, K., Dwight, S.S., Eppig, J.T., et al. (2000). Gene Ontology: tool for the unification of biology. Nature Genetics 2000 25:1 25, 25–29. 10.1038/75556.

110. Wolf, F.A., Angerer, P., and Theis, F.J. (2018). SCANPY: Large-scale single-cell gene expression data analysis. Genome Biol 19, 1–5. 10.1186/S13059-017-1382-0/FIGURES/1.

111. Robinson, J.L., Kocabaş, P., Wang, H., Cholley, P.E., Cook, D., Nilsson, A., Anton, M., Ferreira, R., Domenzain, I., Billa, V., et al. (2020). An Atlas of Human Metabolism. Sci Signal 13, eaaz1482. 10.1126/SCISIGNAL.AAZ1482.

112. Ellis B, H.P.H.F.L.M.N.G.N.S.J.J.M.F.G. (2025). flowCoreL floreCore: Basic structures for flow cytometry data. Preprint.

113. Weber, L.M., Nowicka, M., Soneson, C., and Robinson, M.D. (2019). diffcyt: Differential discovery in high-dimensional cytometry via high-resolution clustering. Commun Biol 2. 10.1038/S42003-019-0415-5.

