## Supplemental Figures for "CD38 expression by neonatal human naïve CD4^+^ T cells shapes their distinct metabolic state and tolerogenic potential"

### Supplementary Figures

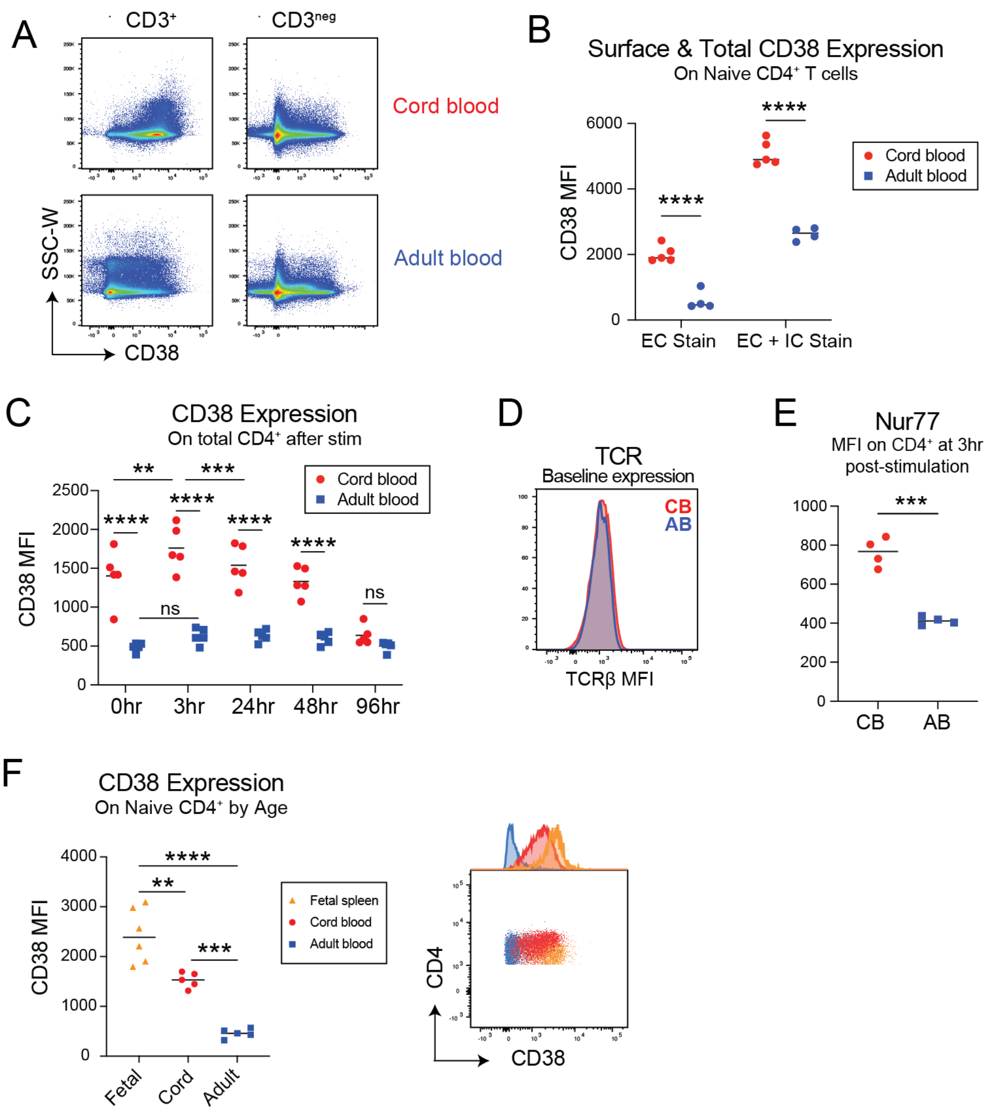


**Figure S1: Related to main Figure 1** (A) Staining and difference of CD38 between CD3+ and CD3- cells from cord blood (CB) and adult blood (AB) at baseline. (B) CD38 MFI on CB and AB naïve CD4+ T cells when stained extracellularly (EC) vs extra- and intracellularly (EC + IC). (C) CD38 MFI on CD4+ T cells at 0, 3, 24, 48, and 96hr after activation with Immunocult$^{TM}$ anti-CD3/CD28 soluble antibodies. (D) TCRβ MFI on naïve CD4+ T cells from CB and AB measured at baseline (0hr). (E) Nur77 mean fluorescence intensity (MFI) in CD4+ T cells measured 3hr post stimulation. (F) CD38 MFI on naïve CD4+ T cells from 21-24 gestational week fetal spleen, CB, or AB.

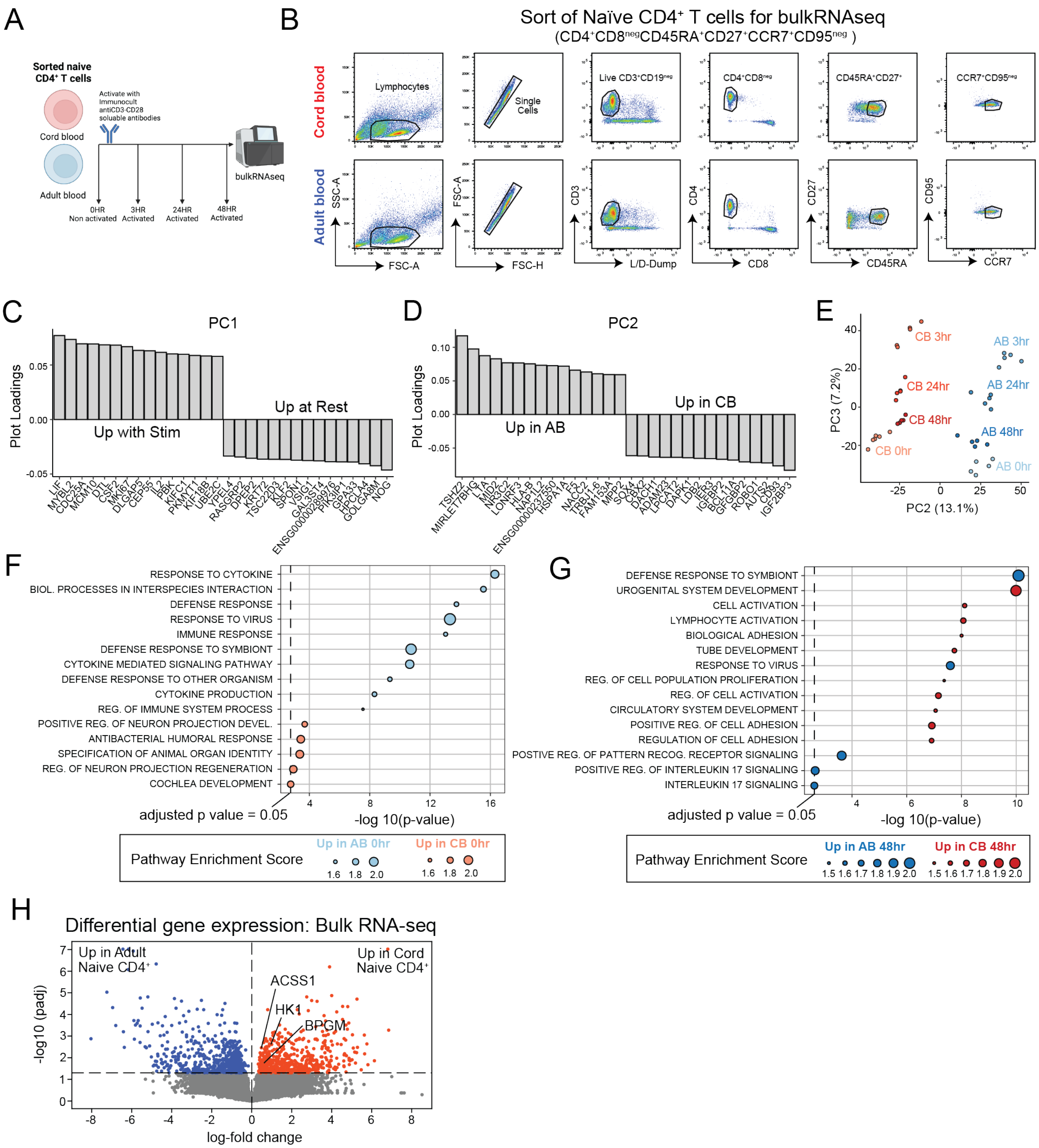


**Figure S2: Related to main Figure 2** (A) Schematic of bulk RNA-seq data acquisition. CB and AB naïve CD4$^+$ T cells were isolated via FACS and sequenced at 0hr (baseline, non-activated) or following 3hr, 24hr or 48hr of activation with ImmunoCult$^{TM}$ anti-CD3/CD28 soluble antibodies. (B) Sort strategy for naïve CD4$^+$ T cells from CB and AB for bulk RNA-seq (Live CD4$^+$CD8$^{neg}$CD45RA$^+$CD27$^+$CCR7$^+$CD95$^{neg}$). (C) Gene loadings with the greatest contributions to PC1 are associated with activation. (D) Gene loadings with the greatest contributions to PC2 are associated with age. (E) PC2 vs. PC3 from bulk RNA-seq dataset. (F-G) Geneset enrichment analysis on ranked genes in CB vs. AB at 0hr and 48hr. (H) Differential expression analysis on bulk RNA-seq data shows upregulation of certain glycolysis-related genes. Genes with statistically significant differential expression are colored by age.

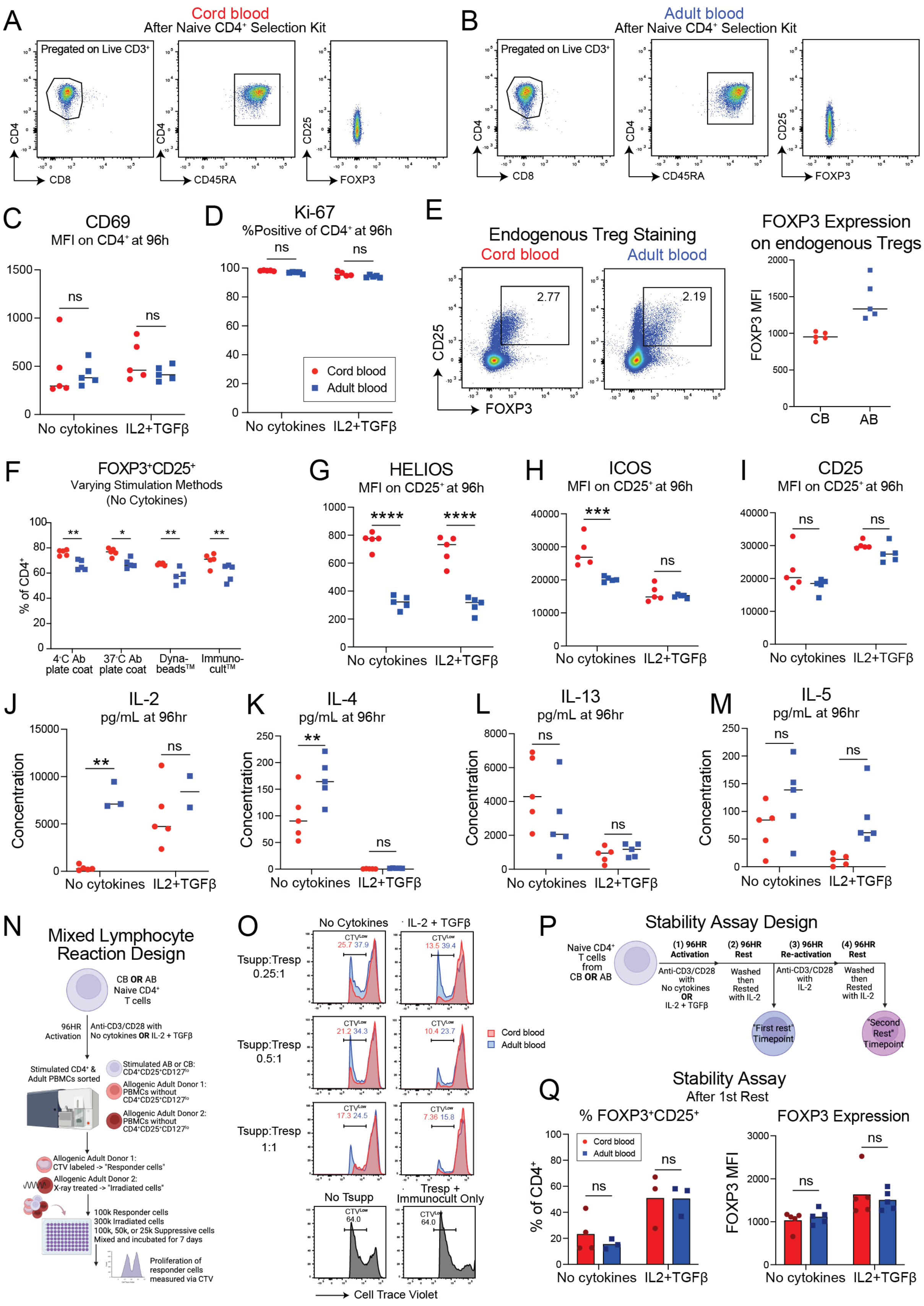
A
Cord blood
After Naive CD4+ Selection Kit
Pregated on Live CD3+
CD4
CD8
CD45RA
CD25
FOXP3
B
Adult blood
After Naive CD4+ Selection Kit
Pregated on Live CD3+
C
CD69
MFI on CD4+ at 96h
ns
No cytokines
IL2+TGFβ
D
Ki-67
%Positive of CD4+ at 96h
Cord blood
Adult blood
E
Endogenous Treg Staining
Cord blood
Adult blood
2.77
2.19
FOXP3 Expression
on endogenous Tregs
FOXP3 MFI
CB
AB
F
FOXP3+CD25+
Varying Stimulation Methods
(No Cytokines)
% of CD4+
4°C Ab plate coat
37°C Ab plate coat
Dyna-beads™
Immuno-cult™
G
HELIOS
MFI on CD25+ at 96h
H
ICOS
MFI on CD25+ at 96h
I
CD25
MFI on CD25+ at 96h
J
IL-2
pg/mL at 96hr
Concentration
K
IL-4
pg/mL at 96hr
L
IL-13
pg/mL at 96hr
M
IL-5
pg/mL at 96hr
N
Mixed Lymphocyte Reaction Design
CB OR AB Naive CD4+ T cells
96HR Activation
Anti-CD3/CD28 with No cytokines OR IL-2 + TGFβ
Stimulated CD4+ & Adult PBMCs sorted
Stimulated AB or CB: CD4+CD25+CD127lo
Allogenic Adult Donor 1: PBMCs without CD4+CD25+CD127lo
Allogenic Adult Donor 2: PBMCs without CD4+CD25+CD127lo
Allogenic Adult Donor 1: CTV labeled -> "Responder cells"
Allogenic Adult Donor 2: X-ray treated -> "Irradiated cells"
100k Responder cells
300k Irradiated cells
100k, 50k, or 25k Suppressive cells
Mixed and incubated for 7 days
Proliferation of responder cells measured via CTV
O
No Cytokines
IL-2 + TGFβ
Tsupp:Tresp 0.25:1
Tsupp:Tresp 0.5:1
Tsupp:Tresp 1:1
CTVLow
25.7 37.9
13.5 39.4
21.2 34.3
10.4 23.7
17.3 24.5
7.36 15.8
Cord blood
Adult blood
No Tsupp
Tresp + Immunocult Only
64.0
Cell Trace Violet
P
Stability Assay Design
Naive CD4+ T cells from CB OR AB
(1) 96HR Activation
Anti-CD3/CD28 with No cytokines OR IL-2 + TGFβ
(2) 96HR Rest
Washed then Rested with IL-2
(3) 96HR Re-activation
Anti-CD3/CD28 with IL-2
(4) 96HR Rest
Washed then Rested with IL-2
"First rest" Timepoint
"Second Rest" Timepoint
Q
Stability Assay
After 1st Rest
% FOXP3+CD25+
% of CD4+
FOXP3 Expression
FOXP3 MFI
Cord blood
Adult blood
No cytokines
IL2+TGFβ

**Figure S3: Related to main Figure 3** (A,B) Representative flow plots showing the EasySep™ Human Naïve CD4$^+$ T Cell Isolation Kit results in a naïve CD4+ T cell population after use from cord blood (CB) and adult blood (AB) peripheral blood mononuclear cells (PBMCs). (C, D) CD69 and Ki67 on total CD4$^+$ population, after naïve CD4$^+$ T cell were isolated from CB and AB and profiled at 96hr after TCR stimulation with Immunocult$^{TM}$ anti-CD3/CD28 soluble antibodies with no added cytokines or with IL-2 (10 ng/ml; PeproTech) and TGF-β (50 ng/ml; PeproTech). (E) Staining patterns for FOXP and CD25 expression on endogenous Tregs from CB and AB whole PBMCs and quantification of FOXP3 MFI on populations gated at left. (F) Percentage of FOXP3$^+$CD25$^+$ cells from CB and AB naïve CD4$^+$ T cells at 96hr after TCR stimulation by different methods, including culture in a plate coated with anti-CD3/CD28 antibodies overnight at 4°C, in a plate coated with anti-CD3/CD28 antibodies for 4hr at 37°C, with beads added to culture coated in anti-CD3/CD28 (Dynabeads™), or with Immunocult™ anti-CD3/CD28 soluble antibodies. All with no added cytokines. (G-I) Expression of various markers on FOXP3$^+$CD25$^+$ at 96hr after CB or AB naïve CD4$^+$ T cell stimulation, as measured by mean fluorescence intensity (MFI) by flow cytometry. (J-M) ELISA-based measurement of cytokines in cell culture supernatant of stimulated CB and AB naïve CD4$^+$ T cells at 96hr. (N) Schematic of mixed lymphocyte reaction ("Allo-MLR") where CB and AB naïve CD4+ T cells were activated with Immunocult$^{TM}$ anti-CD3/CD28 soluble antibodies and cultured for 96hr under no cytokine or IL-2+TGF-β conditions. Cells were then sorted to obtain CD4$^+$CD25$^+$CD127$^{LO}$ cells, "Suppressor cells". Two unrelated AB donors' PBMCs were thawed and sorted to remove CD4$^+$CD25$^+$CD127$^{LO}$ cells from those samples. One was stained with CTV ("Responder cells") while the other was irradiated ("Irradiated cells") to serve as a source of antigen-presenting cells. Cells were then mixed together at the ratio of 100,000 Responder cells, 300,000 Irradiated cells, and 100,000, 50,000, or 25,000 Suppressive cells with no added anti-CD3/CD28 stimulation. Proliferation was measured via CTV after 7 days of culture. (O) Cell Trace Violet (CTV) expression from Allo-MLR cultures showing proliferation levels of Responder cells between conditions. Responder cells + Irradiated cells (No Tsupp) as well as Responder cells with ImmunoCult and no Suppressive or Irradiated cells were used as controls for proliferation of Responder cells. (P) Schematic of stability assay. FOXP3 and CD25 expression stability was assessed in CB and AB where naïve CD4$^+$ T cells were activated with ImmunocultTM anti-CD3/CD28 soluble antibodies and cultured for 96hr under no cytokine or IL-2+TGF-β conditions. Cells were then washed to remove stimulation and cytokines and rested with 96hr with IL-2 in all conditions. After the first 96hr rest, half the cells were removed to assess FOXP3 and CD25 expression. Remaining cells were then re-stimulated with ImmunocultTM anti-CD3/CD28 soluble antibodies and IL-2 in all conditions for 96hr. Following the second stimulation, cells were washed and rested with IL-2 for 96hr. After the second 96hr rest, FOXP3 and CD25 expression were assessed. (Q) FOXP$^+$CD25$^+$ expression after the first rest in the stability assay.

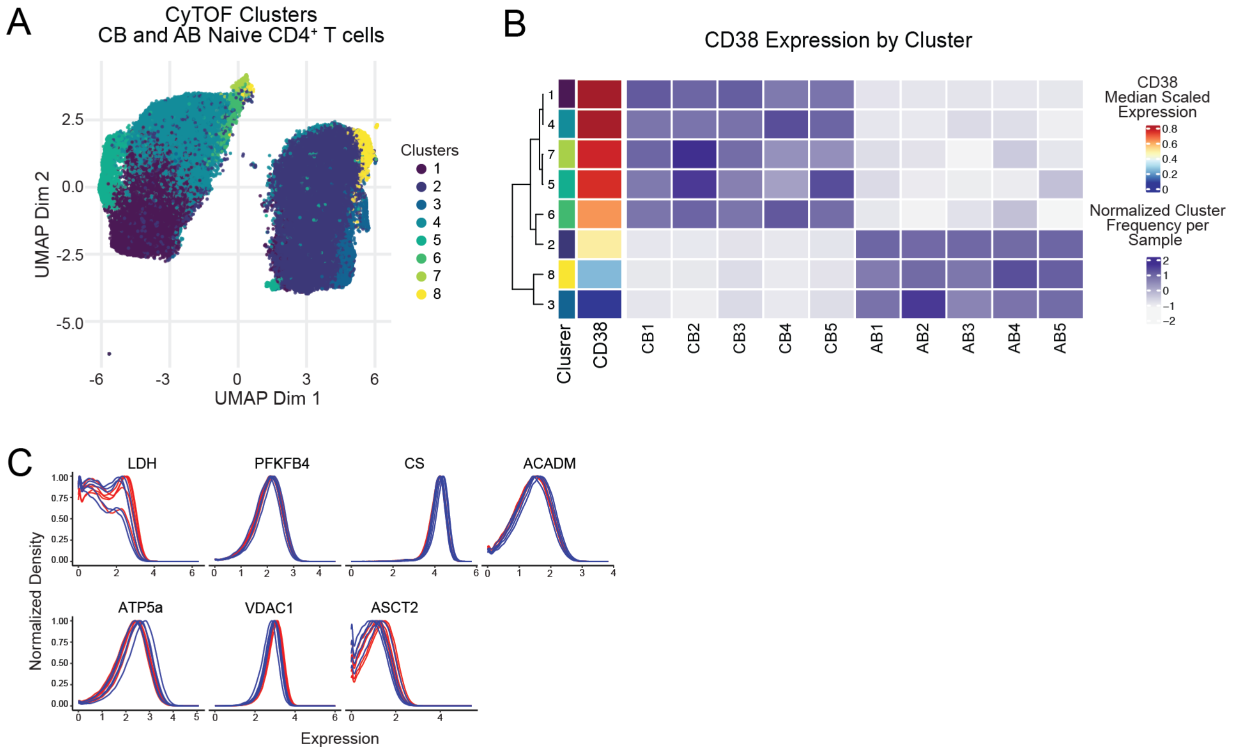


**Figure S4: Related to main Figure 4** (A) To examine the different metabolic states present in the naïve CD4+ T cell pool across ages, clustering was performed with the FlowSOM algorithm using metabolic and reduced lineage markers, yielding 8 distinct clusters. (B) CD38 expression across the 8 distinct clusters in CB and AB naïve CD4+ T cells. (C) Expression patterns of metabolic markers for all CB and AB donors.

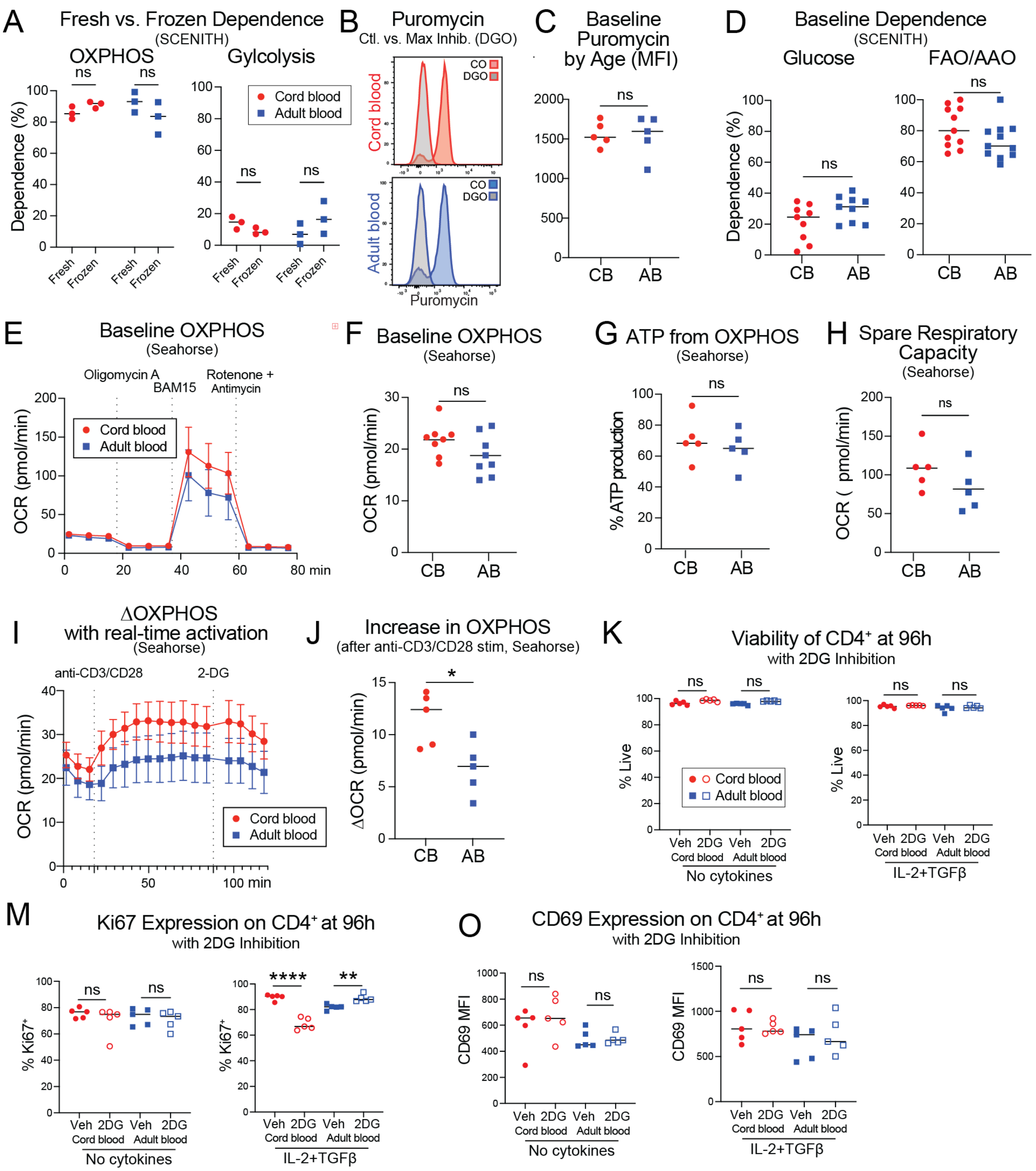


**Figure S5: Related to main Figure 5** (A) Baseline (non-activated) dependence on glycolysis and mitochondrial oxidative phosphorylation in fresh and previously frozen cord and adult CD4+ T cells measured via SCENITH. (B) Example staining for SCENITH of puromycin MFI shift with Control vs DGO conditions for baseline (non-activated) CB and AB naive CD4+ T cells. (C) Quantification of puromycin MFI in Control (no metabolic inhibitor condition) for baseline (non-activated) CB and AB naive CD4+ T cells. (D) Baseline (non-activated) dependence on glucose and fatty acid/amino acid oxidation in CB and AB naive CD4+ T cells measured via SCENITH. (E) Seahorse graph of OCR (oxidative phosphorylation (OXPHOS)) of baseline (non-activated) CB and AB naive CD4+ T cells. (F) Quantification of baseline OXPHOS between CB and AB naive CD4+ T cells via Seahorse. (G) The percent of ATP made from OXPHOS between CB and AB naive CD4+ T cells via Seahorse. (H) Spare respirator capacity between CB and AB naive CD4+ T cells via Seahorse. (I) Seahorse graph of ECAR after acute injection with anti-CD3/CD28 soluble antibodies between CB and AB naive CD4+ T cells via Seahorse. (J) The change in OCR after acute injection with anti-CD3/CD28 soluble antibodies between CB and AB naive CD4⁺ T cells via Seahorse. (K-O) Quantification of total live cells (K), Ki67⁺ (M), and CD69 MFI (O) on CD4+ T cells after 24 hours of treatment with 0.5mM 2DG followed by activation with anti-CD3/CD28 soluble antibodies with no added cytokines or with IL-2 + TGF-β, measured 96 hours after activation.

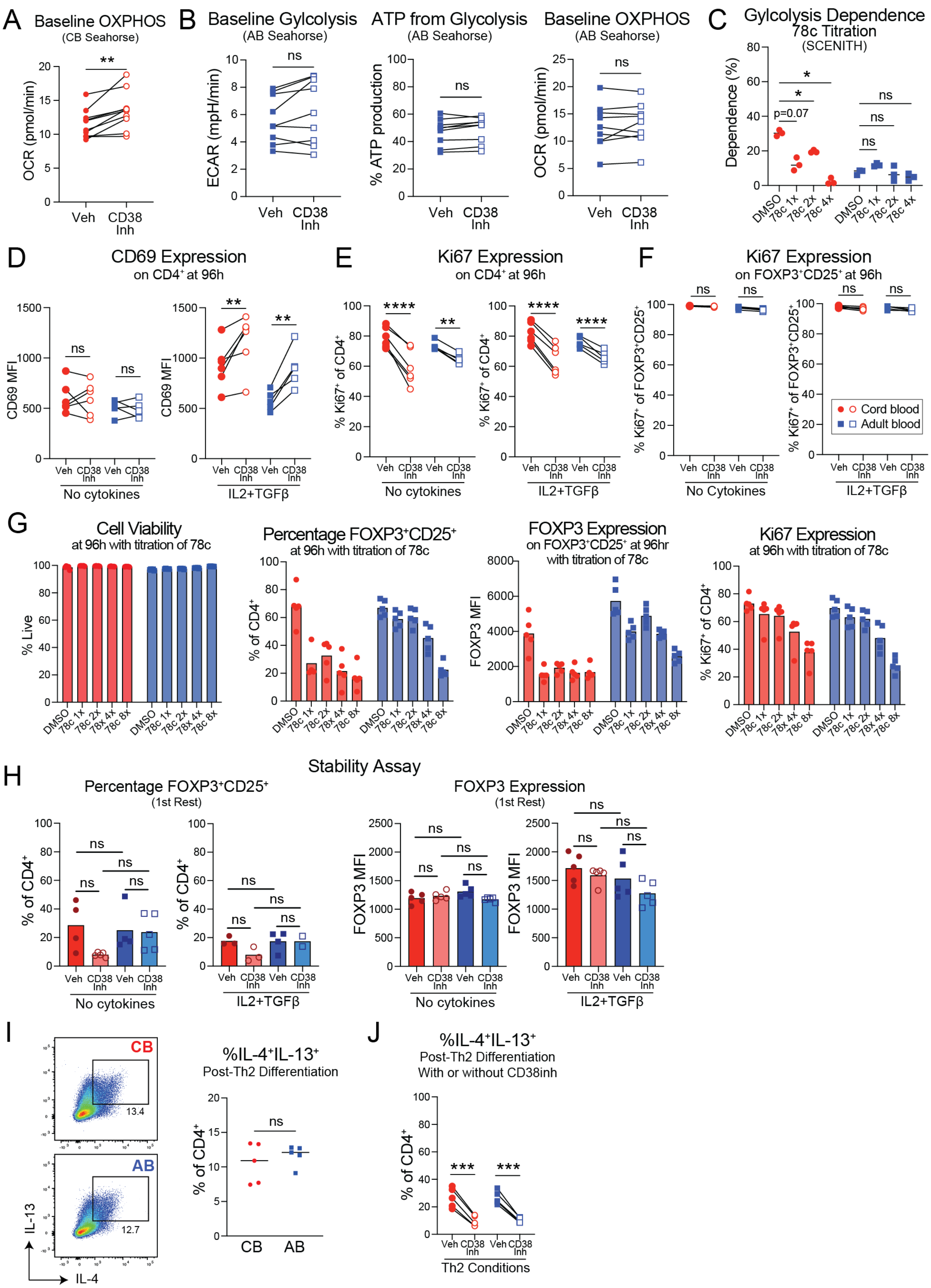
A
Baseline OXPHOS
(CB Seahorse)
OCR (pmol/min)
Veh
CD38 Inh
B
Baseline Gylcolysis
(AB Seahorse)
ECAR (mpH/min)
ATP from Glycolysis
(AB Seahorse)
% ATP production
Baseline OXPHOS
(AB Seahorse)
OCR (pmol/min)
C
Gylcolysis Dependence
78c Titration
(SCENITH)
Dependence (%)
p=0.07
DMSO
78c 1x
78c 2x
78c 4x
D
CD69 Expression
on CD4+ at 96h
CD69 MFI
No cytokines
IL2+TGFβ
E
Ki67 Expression
on CD4+ at 96h
% Ki67+ of CD4+
F
Ki67 Expression
on FOXP3+CD25+ at 96h
% Ki67+ of FOXP3+CD25+
No Cytokines
Cord blood
Adult blood
G
Cell Viability
at 96h with titration of 78c
% Live
Percentage FOXP3+CD25+
at 96h with titration of 78c
% of CD4+
FOXP3 Expression
on FOXP3+CD25+ at 96hr
with titration of 78c
FOXP3 MFI
Ki67 Expression
at 96h with titration of 78c
% Ki67+ of CD4+
78c 8x
H
Stability Assay
Percentage FOXP3+CD25+
(1st Rest)
FOXP3 Expression
(1st Rest)
I
CB
13.4
AB
12.7
IL-13
IL-4
%IL-4+IL-13+
Post-Th2 Differentiation
J
%IL-4+IL-13+
Post-Th2 Differentiation
With or without CD38inh
Th2 Conditions

**Figure S6: Related to main Figure 6** (A) (Baseline (non-activated) OXPHOS in CB naïve CD4$^+$ T cells after 24hr of 5uM 78c treatment measured via Seahorse. (B) Baseline (non-activated) glycolysis, ATP from glycolysis, and OXPHOS in AB naïve CD4$^+$ T cells after 24hr of 78c treatment measured via Seahorse. (C) Glycolytic dependence measured via SCENITH in CB and AB naïve CD4$^+$ T cells after 24 hours of treatment with various doses of 78c (1x= 5uM). (D-F) Quantification at 96 hour post-stimulation of CD69 MFI on total CD4$^+$ (D), Ki67$^+$ on total CD4$^+$ (E), and Ki67$^+$ on FOXP3$^+$CD25$^+$ (F) after 24 hours of pre-treatment with 5uM 78c followed by "No Cytokine" or "IL-2 + TGF-β" activation in the ongoing presence of 78c. (G) Cell viability, percent FOXP3$^+$CD25$^+$, FOXP3 MFI on FOXP3$^+$CD25$^+$, and percent Ki67$^+$ in the setting of "No Cytokine" stimulation with various doses of 78c (1x = 5uM). (H) FOXP3$^+$CD25$^+$ and FOXP3 expression from first rest of stability assay with 5uM 78c added for 24hr before activation and during the first activation. (I) IL-4 and IL-13 expression was measured from Th2 differentiation of naïve CD4$^+$ T cells from CB and AB. (J) IL-4 and IL-13 expression was measured from Th2 differentiation of naïve CD4$^+$ T cells from CB and AB where 5uM 78c was added for 24hr prior to and during the Th2 differentiation.

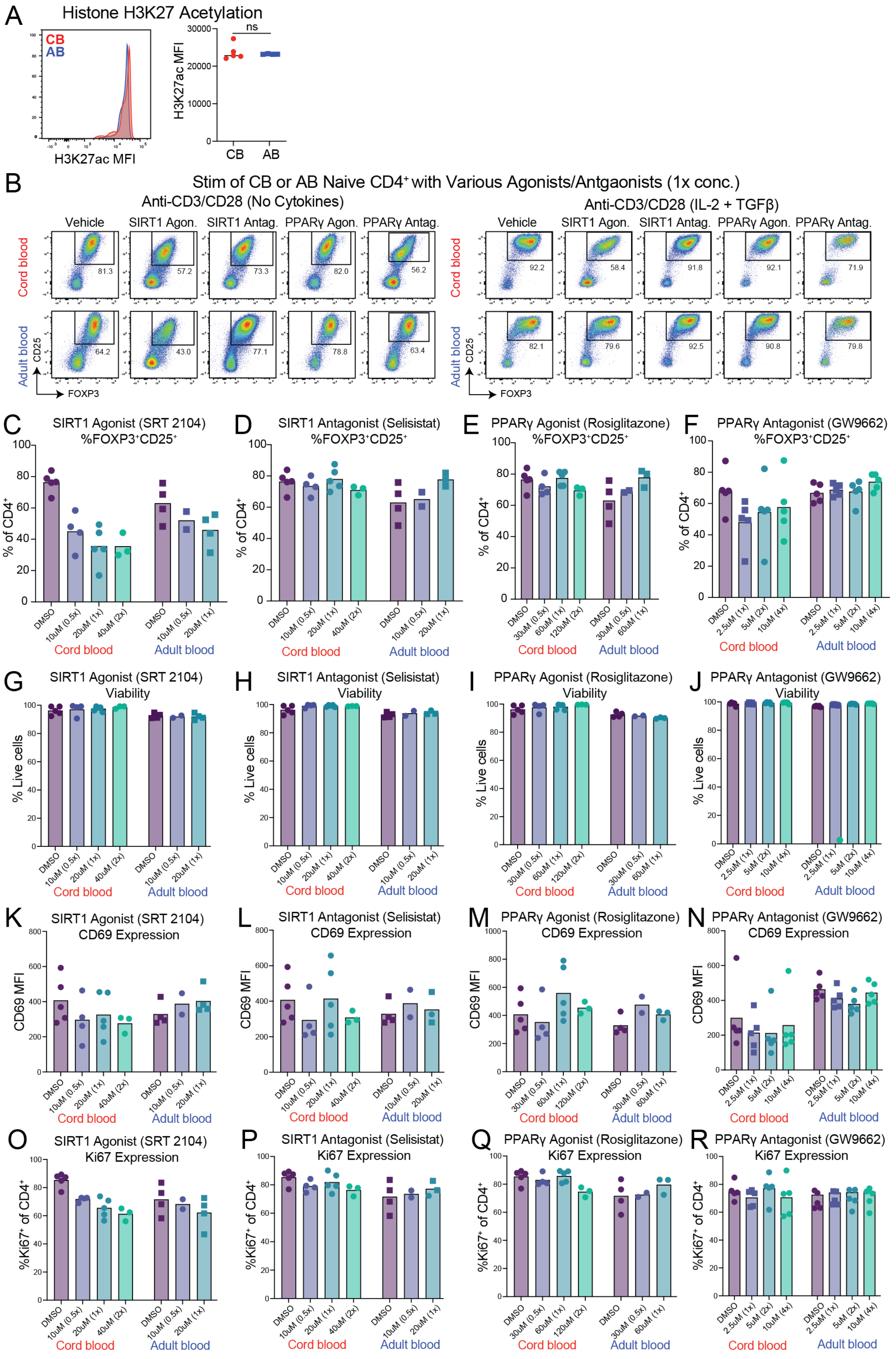
A
Histone H3K27 Acetylation
CB
AB
H3K27ac MFI
ns
H3K27ac MFI
CB
AB
B
Stim of CB or AB Naive CD4+ with Various Agonists/Antgaonists (1x conc.)
Anti-CD3/CD28 (No Cytokines)
Anti-CD3/CD28 (IL-2 + TGFβ)
Vehicle
SIRT1 Agon.
SIRT1 Antag.
PPARγ Agon.
PPARγ Antag.
Cord blood
Adult blood
CD25
FOXP3
C
SIRT1 Agonist (SRT 2104) %FOXP3+CD25+
D
SIRT1 Antagonist (Selisistat) %FOXP3+CD25+
E
PPARγ Agonist (Rosiglitazone) %FOXP3+CD25+
F
PPARγ Antagonist (GW9662) %FOXP3+CD25+
% of CD4+
G
SIRT1 Agonist (SRT 2104) Viability
H
SIRT1 Antagonist (Selisistat) Viability
I
PPARγ Agonist (Rosiglitazone) Viability
J
PPARγ Antagonist (GW9662) Viability
% Live cells
K
SIRT1 Agonist (SRT 2104) CD69 Expression
L
SIRT1 Antagonist (Selisistat) CD69 Expression
M
PPARγ Agonist (Rosiglitazone) CD69 Expression
N
PPARγ Antagonist (GW9662) CD69 Expression
CD69 MFI
O
SIRT1 Agonist (SRT 2104) Ki67 Expression
P
SIRT1 Antagonist (Selisistat) Ki67 Expression
Q
PPARγ Agonist (Rosiglitazone) Ki67 Expression
R
PPARγ Antagonist (GW9662) Ki67 Expression
%Ki67+ of CD4+
DMSO
10uM (0.5x)
20uM (1x)
40uM (2x)
30uM (0.5x)
60uM (1x)
120uM (2x)
2.5uM (1x)
5uM (2x)
10uM (4x)
Cord blood
Adult blood

**Figure S7: Related to main Figure 7** (A) H3K27ac staining of CB and AB naïve $CD4^+$ T cells using an antibody against Acetyl-Histone H3 (Lys27) (Cell Signaling Tech, # 39030S). (B) Flow plots of $FOXP3^+CD25^+$ cells from CB and AB "No Cytokine" and "IL-2 + TGF-β" culture conditions treated with DMSO (control) or pharmacological compounds. (C-R) Quantification on $CD4^+$ cells of $FOXP3^+CD25^+$ percentages (C-F), cell viability (G-J), activation as marked by CD69 MFI (K-N) or recent proliferation as indicated by percentage of $Ki67^+$ (O-R) from CB and AB "No Cytokine" culture conditions treated with DMSO (vehicle control) or various doses of several pharmacological compounds.
